# The antibiotic phazolicin displays a dual mode of uptake in Gram-negative bacteria

**DOI:** 10.1101/2022.04.27.489825

**Authors:** Dmitrii Y. Travin, Armelle Vigouroux, Satomi Inaba-Inoue, Feng Qu, Romain Jouan, Joy Lachat, Dmitry Sutormin, Svetlana Dubiley, Konstantinos Beis, Solange Moréra, Konstantin Severinov, Peter Mergaert

**Author notes:** To whom correspondence should be addressed (K.S.); (P.M.). J.L.: Center for Immunology and Infectious Diseases, Cimi-Paris, Inserm, Sorbonne Université, Paris, France.

## Abstract

Phazolicin (PHZ) is a peptide antibiotic exhibiting narrow-spectrum activity against rhizobia closely related to its producer *Rhizobium* sp. Pop5. Using genetic and biochemical techniques, we here identified BacA and YejABEF as two importers of PHZ in a sensitive model strain *Sinorhizobium meliloti* Sm1021. BacA and YejABEF are members of SLiPT and ABC transporter families of non-specific peptide importers, respectively. The uptake of PHZ by two distinct families of transporters dramatically decreases the naturally occurring rate of resistance. Moreover, since both BacA and YejABEF are essential for the development of functional symbiosis of rhizobia with leguminous plants, the acquisition of PHZ resistance via the inactivation of transporters is further disfavoured since single *bacA* or *yejABEF* mutants are unable to propagate in root nodules. Crystal structures of the periplasmic subunit YejA from *S. meliloti* and *Escherichia coli* revealed fortuitous bound peptides, suggesting a non-specific peptide-binding mechanism that facilitates the uptake of PHZ and other antimicrobial peptides.

**SIGNIFICANCE:** Many bacteria produce antimicrobial peptides to eliminate competitors and create an exclusive niche. These peptides kill bacteria by either membrane disruption or inhibiting essential intracellular processes. The Achilles heel of the latter type of antimicrobials is their dependence on transporters to enter the susceptible bacteria since mutations in such transporters result in resistance. We describe here how the ribosome-targeting peptide phazolicin, produced by *Rhizobium* sp. Pop5, uses two different transporters, BacA and YejABEF, to get into the cells of the symbiotic bacterium *Sinorhizobium meliloti*. This dramatically reduces the probability of resistance acquisition. Both transporters need to be inactivated for phazolicin resistance acquisition. Since these transporters are also crucial in *S. meliloti* for its symbiotic association with host plants, their inactivation in biological settings is highly unlikely. This makes PHZ an attractive lead for the development of a biocontrol agent with potential for use in agriculture.

## INTRODUCTION

Molecules of peptidic nature constitute a considerable part of the known diversity of naturally occurring compounds with antimicrobial activity [1]. The passage of intracellularly acting peptidic antibiotics through the cell envelope is a step limiting their effective concentration [2,3]. Many of these compounds rely on non-specific peptide transporters for internalization. Since mutations affecting antibiotic uptake constitute one of the major sources of resistance development to antimicrobial peptides [4], understanding the transport pathways for newly identified molecules is an important objective.

Phazolicin (PHZ) is a recently discovered azole-containing peptide produced by soil bacterium *Rhizobium* sp. Pop5 [5]. PHZ belongs to a class of natural products known as <u>ri</u>bosomally-synthesized and <u>p</u>osttranslationally-modified <u>p</u>eptides (RiPPs) [6,7]. It contains eight azole cycles installed into the structure of a precursor peptide through the posttranslational cyclization of Ser and Cys side chains by specific enzymes (**Fig. 1A**). PHZ inhibits, at low micromolar concentrations, the growth of *Rhizobium* and *Sinorhizobium* strains closely related to its producer. PHZ is a translation inhibitor acting through the obstruction of the peptide exit channel of the 50S ribosome subunit of susceptible cells [5]. The PHZ-producing *Rhizobium* sp. Pop5 is a nitrogen-fixing bacterium that can establish symbiotic relationships with common beans *Phaseolus vulgaris* through the formation and infection of nodules – specific symbiotic root organs. Production of an unrelated RiPP, trifolitoxin, was shown to enhance the competitiveness of its producer *Rhizobium leguminosarum* bv. *trifolii* T24 in root colonization and the occupancy of nodules [8,9]. The increased competitiveness of trifolitoxin producer is likely due to its ability to inhibit the growth of related strains capable of plant colonization. By analogy, PHZ may also enable *Rhizobium* sp. Pop5 to outcompete other strains in either soil or host plant niches (or both).

**Figure 1.**
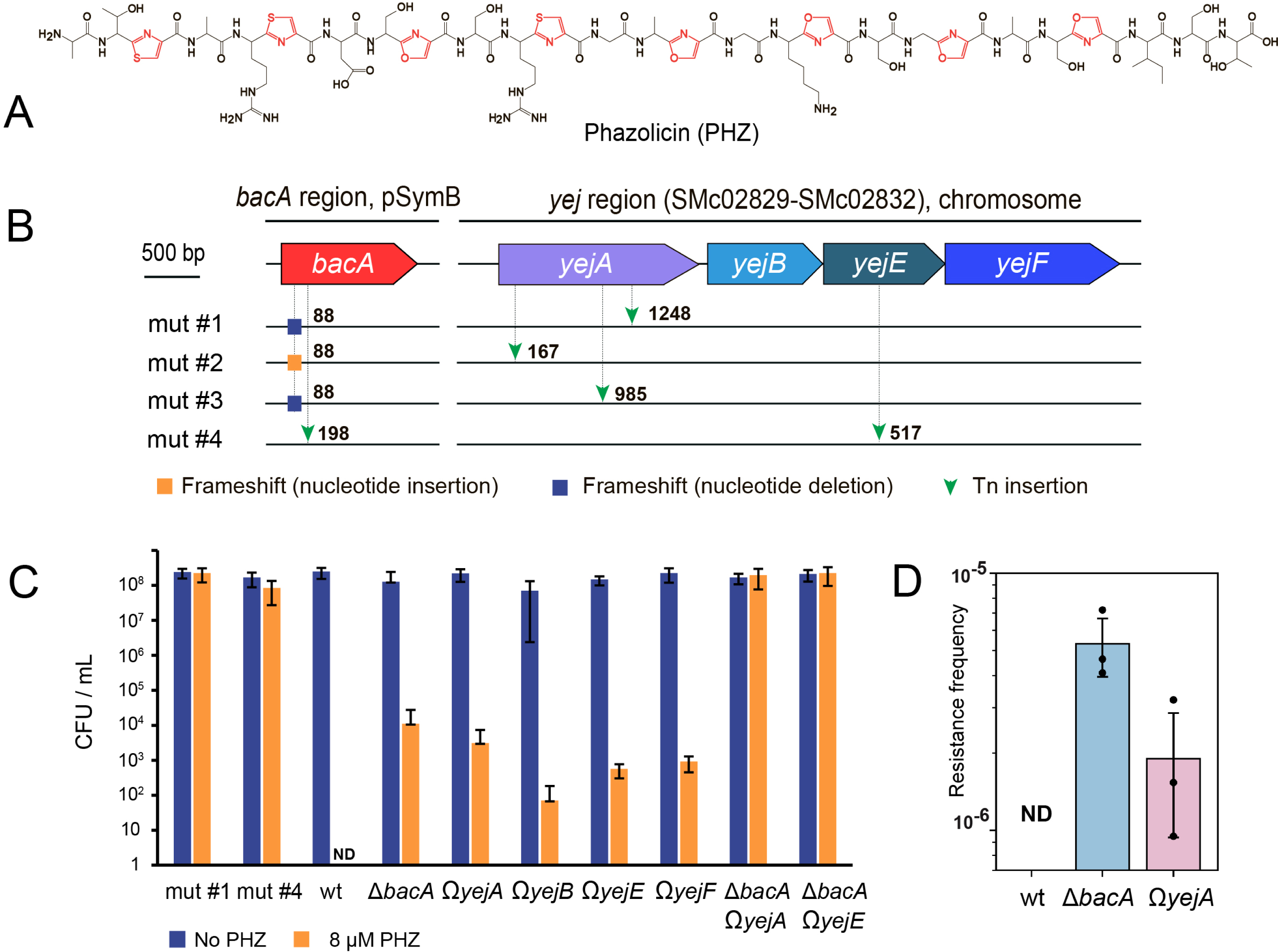
BacA and YejABEF transporters independently contribute to the uptake of phazolicin by *Sinorhizobium meliloti*. **(A)** Chemical structure of phazolicin. Post-translationally installed thiazole and oxazole cycles are shown in red. **(B)** Schematic representation of mutations identified by whole-genome sequencing of four PHZ-resistant mutants of Sm1021. The numbers indicate the nucleotide position in the gene, where the mutation was detected. **(C)** CFU of Sm1021 growing on the medium with (8 μM) and without PHZ. Results are presented as the average of three biological replicates for obtained PHZ-resistant strains (mut #1 and mut #4), single and double mutants in *bacA* and *yejABEF* genes. ND – not detected. **Fig. S1** shows a picture of the CFU plating for one replicate. **(D)** Frequency of PHZ resistance acquisition in *wt* Sm1021, *bacA* and *yejA* single gene mutants. The data for three biological replicates are shown. ND, not detected.

Here, we show that peptide transporters from two different families, the ATP-binding cassette (ABC) importer YejABEF and the SbmA-like peptide transporter (SLiPT) BacA, independently import PHZ into the cells of *Sinorhizobium meliloti*. BacA is a dimer composed of two identical membrane-spanning subunits capable of peptide/H^+^ symport [10]. BacA of *S. meliloti* was previously shown to internalize antimicrobial peptides including bleomycin (BLM) – a hybrid peptide-polyketide antibiotic acting as a DNA-damaging agent [11]. SbmA, a BacA homolog from *E. coli*, is responsible for the uptake of such chemically and functionally diverse compounds as RNA polymerase-targeting lasso-peptide microcin J25 [12], gyrase inhibitor microcin B17 [13], BLM [14], and ribosome-targeting peptides Bac7 [15] and klebsazolicin [16]. While the structure of YejABEF is unknown, based on similarity with other ABC-transporters, it is most likely composed of two transmembrane subunits (YejB and YejE), two nucleotide-binding subunits (YejF_2_), and a periplasmic substrate-binding subunit (YejA). In *E. coli*, YejABEF was previously shown to be the entry point of microcin C, a peptidyl-nucleotide antibiotic targeting aspartyl-tRNA synthetase [17]. Microcin C exploits its peptide part to get into the cells of susceptible bacteria *via* YejABEF, which is followed by the release of the nucleotide-containing toxic moiety after proteolytic cleavage of the peptide part in the cytoplasm (the so-called Trojan horse mechanism, reviewed in [18]).

In this work, we apply genetic, biochemical, structural, and microbiological approaches to characterize the uptake of PHZ by *S. meliloti*. We further make use of the transporter-deficient PHZ-resistant mutants to demonstrate that PHZ biosynthesis enables its producer to eliminate susceptible bacteria and create a selective niche.

## RESULTS

### *PHZ-resistant* S. meliloti *mutants simultaneously carry mutations in genes of two independent peptide import systems*

To identify the transporters mediating the internalization of phazolicin into the cells of PHZ-susceptible *S. meliloti* Sm1021, we attempted to select spontaneous resistant mutants by plating aliquots of overnight cultures on solid medium containing 20 μM PHZ. However, no resistant mutants were recovered, a surprising result given that the same approach readily yielded *E. coli* mutants resistant to azole-modified peptide microcin B17 with lesions in the gene of its transporter [13].

Since the direct approach to the isolation of PHZ-resistant strains was unsuccessful, we constructed a Mariner Himar C9 transposon library of *S. meliloti* Sm1021 and screened it on PHZ-containing solid medium as described above. In this way, four colonies growing on the medium with 20 μM PHZ were obtained. To identify the mutations they carried, whole-genome Illumina sequencing of cells from each colony was performed. The BLAST search for the transposon sequence in the NGS data identified transposon insertions in the *bacA, yejA*, and *yejE* genes (**Fig. 1B**). The *bacA* gene, located on the 1.68 Mbp pSymB *S. meliloti* Sm1021 megaplasmid, encodes a peptide transporter homologous to the *E. coli* SbmA [10]. The chromosomally-located *yejA* and *yejE* encode, respectively, the periplasmic component and the permease of the YejABEF ABC importer [19]. Noteworthy, mutants #1, #2, and #3, which had transposon insertions in *yejA*, also carried point mutations shifting the reading frame at the beginning of the *bacA* gene. In mutant #4, there was a transposon insertion in the *bacA* gene. In addition, at the nucleotide 517 of the *yejE* gene, sequences of transposon ends were detected. Thus, all four selected PHZ-resistant lineages turned out to be double mutants with both the BacA and the YejABEF inner membrane uptake systems inactivated. This suggests that the simultaneous occurrence of two mutations is necessary to acquire PHZ resistance and explains our initial failure to select resistant mutants in the *wt* Sm1021.

### BacA and YejABEF independently contribute to the uptake of phazolicin

To confirm the role of BacA and YejABEF transporters in the uptake of PHZ, we plated serial dilutions of cultures of the previously constructed Sm1021 deletion strain Δ*bacA* [20] and of four plasmid insertion mutants in the *yejA, yejB, yejE*, and *yejF* genes (designated Ω*yejA* through Ω*yejF*) on plates supplemented with PHZ. *Wt* Sm1021 and mutants #1 and #4 served as negative (susceptible) and positive (resistant) controls, respectively. As can be seen in **Figs. 1C and S1**, single mutants in either *bacA* or any of the *yej* genes were susceptible to phazolicin. However, in contrast to the *wt* we observed the growth of separate colonies where aliquots of undiluted or 10-fold diluted cultures of single mutants were applied on PHZ-containing plates (**Fig. S1**). Mutants #1 and #4 were fully resistant to PHZ as expected. Together, these results suggest that the presence of either one of the two transport systems is sufficient to import enough PHZ for nearly full toxicity and that both transporters need to be inactivated to confer resistance.

To rule out the possibility that additional mutations contributing to PHZ-resistance have accumulated in genes other than *bacA* and *yejABEF* during transposon library construction and screening, a double mutant devoid of both transporter systems was constructed. We used phiM12 phage lysate of Sm1021 Δ*bacA* (Sp^R^) to transduce the *bacA* mutation into the Ω*yejA* or Ω*yejE* (Km^R^) mutants. The absence of functional *bacA* in transductants selected on Km and Sp was confirmed by PCR. Δ*bacA* Ω*yejA* and Δ*bacA* Ω*yejE* double mutants were as resistant to PHZ as transposon mutants #1 and #4 (**Fig. 1C, Fig. S1**, and **Table 1)**. These data demonstrate that BacA and YejABEF transporters independently contribute to the uptake of PHZ by the cells of *S. meliloti* Sm1021. No other transporters appear to be involved in PHZ transport, at least in our conditions.

**Table 1.**
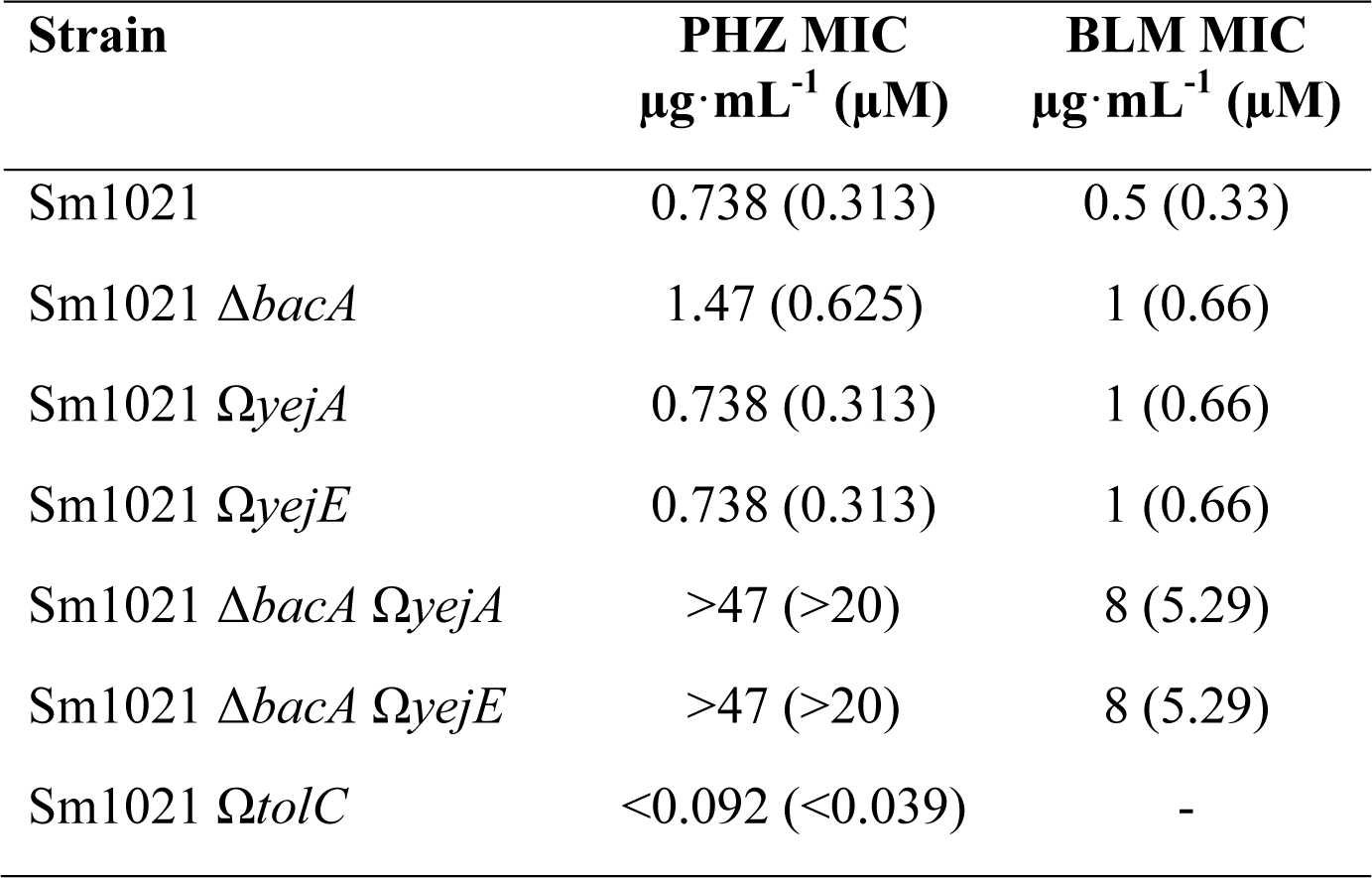
Phazolicin (PHZ) and bleomycin (BLM) MIC values for Sm1021 and its derivatives.

| Strain | PHZ MIC<br>$\mu\text{g}\cdot\text{mL}^{-1}$ ($\mu\text{M}$) | BLM MIC<br>$\mu\text{g}\cdot\text{mL}^{-1}$ ($\mu\text{M}$) |
| --- | --- | --- |
| Sm1021 | 0.738 (0.313) | 0.5 (0.33) |
| Sm1021 $\Delta bacA$ | 1.47 (0.625) | 1 (0.66) |
| Sm1021 $\Omega yejA$ | 0.738 (0.313) | 1 (0.66) |
| Sm1021 $\Omega yejE$ | 0.738 (0.313) | 1 (0.66) |
| Sm1021 $\Delta bacA \Omega yejA$ | >47 (>20) | 8 (5.29) |
| Sm1021 $\Delta bacA \Omega yejE$ | >47 (>20) | 8 (5.29) |
| Sm1021 $\Omega tolC$ | <0.092 (<0.039) | - |

The growth of separate colonies derived from single-gene mutants in the 0 and 10x dilutions (**Fig. S1)** could indicate that these single mutants are slightly less susceptible to PHZ than the *wt* or that spontaneous fully resistant double mutants appear on the single-mutant background. We compared the frequency of spontaneous resistance acquisition in the Δ*bacA* and the Ω*yejA* single mutants and the *wt* Sm1021. While no resistant clones were obtained for the *wt* as above, the rate of resistance acquisition for single mutants was between 10^−5^ and 10^−6^ (**Fig. 1D**). To understand the genetic basis of resistance acquired on the background of inactivated *yejA*, we picked up, at random, six resistant clones of the Sm1021 Ω*yejA* strain (two from each biological replicate) and amplified their *bacA* genomic regions with specific primers. Sanger sequencing revealed mutations in 5 out of 6 *bacA* amplicons (**Fig. S2A**). In three cases, single nucleotide insertions or deletions led to premature stop codon formation in the *bacA* reading frame. Two other mutations led to single amino acid substitutions (L158R and F162S) in one of the BacA transmembrane α-helixes. Since Leu^158^ and Phe^162^ face the inner part of the membrane (**Fig. S2B** and **S2C**) substituting these residues with charged or polar ones should inactivate the transporter. In the remaining mutant, no mutation in the *bacA* open reading frame was detected and we speculate that a mutation in the promoter or another *bacA* regulatory element could inactivate BacA synthesis in this clone. Overall, this data confirms that mutations in *bacA* are the primary source of acquiring resistance by the strain with inactivated YejABEF. We assume that the complementary result would be obtained in PHZ resistant mutants selected on the *bacA* mutant background; we did not check this conjecture experimentally, since the large size of the *yejABEF* operon complicates identification of second-site mutations.

### *PHZ-resistant Sm1021 avoids elimination by* Rhizobium *sp. Pop5 in co-culture*

We were interested to determine whether the resistance of the Sm1021 double Δ*bacA* Ω*yejA* mutant to PHZ could help this strain to escape elimination by the phazolicin producer during co-cultivation. To make sure that any effect on Sm1021 derivatives growth observed can be unambiguously linked to the PHZ production by *Rhizobium* sp. Pop5, a non-producing derivative was constructed and used as a control (**Fig. S3A, B**). This mutant, Pop5 Ω*phzD*, has a plasmid insertion in the *phzD* gene coding for the YcaO domain-containing cyclodehydratase predicted to catalyze PHZ posttranslational modifications [5], [21]. MALDI-ToF mass spectrometry confirmed that *Rhizobium* sp. Pop5 Ω*phzD* was not producing mature PHZ (**Fig. S3C**). Consistently, no growth inhibition zones around the colonies of *Rhizobium* sp. Pop5 Ω*phzD* spotted over a lawn of the PHZ-susceptible *R. leguminosarum* 4292 were observed (**Fig. S3D**). It is worth noting that the phenotype of *Rhizobium* sp. Pop5 Ω*phzD* experimentally confirms the association of PHZ with its biosynthetic gene cluster (BGC), which up to now was based on the sequence of the precursor peptide only [5].

We used flow cytometry to quantify GFP-marked *S. meliloti* derivatives (Sm1021 *wt* expressing GFP encoded on the plasmid pDG71 [22] and transporter-deficient Sm1021 Δ*bacA* Ω*yejA* expressing GFP from the plasmid insertion in the *yejA* gene) and DsRed-marked *Rhizobium* sp. Pop5 derivatives (Pop5 *wt* and Pop5 Ω*phzD* expressing DsRed from plasmid pIN72) after co-cultivation (**Fig. 2A, Fig. S4A**). The analysis at the onset of the co-cultivation (t = 0h) showed a roughly 1:1 ratio of *Rhizobium* and *Sinorhizobium* in the four tested Sm1021-derivative:Pop5-derivative mixtures. After growth for 24h, 48h, or 88h, the Sm1021 *wt* population was strongly reduced in cultures containing PHZ-producing Pop5 *wt*. Moreover, the GFP fluorescence level of the Sm1021 cells remaining in co-cultures diminished compared to the fluorescence level of cells in the initial mixture, indicating that GFP synthesis in Sm1021 is inhibited upon co-cultivation with the PHZ producer, as expected. In other co-cultivation combinations, the proportion of Sm1021 did not drop or even increased and the GFP fluorescence of cells remained at initial levels.

**Figure 2.**
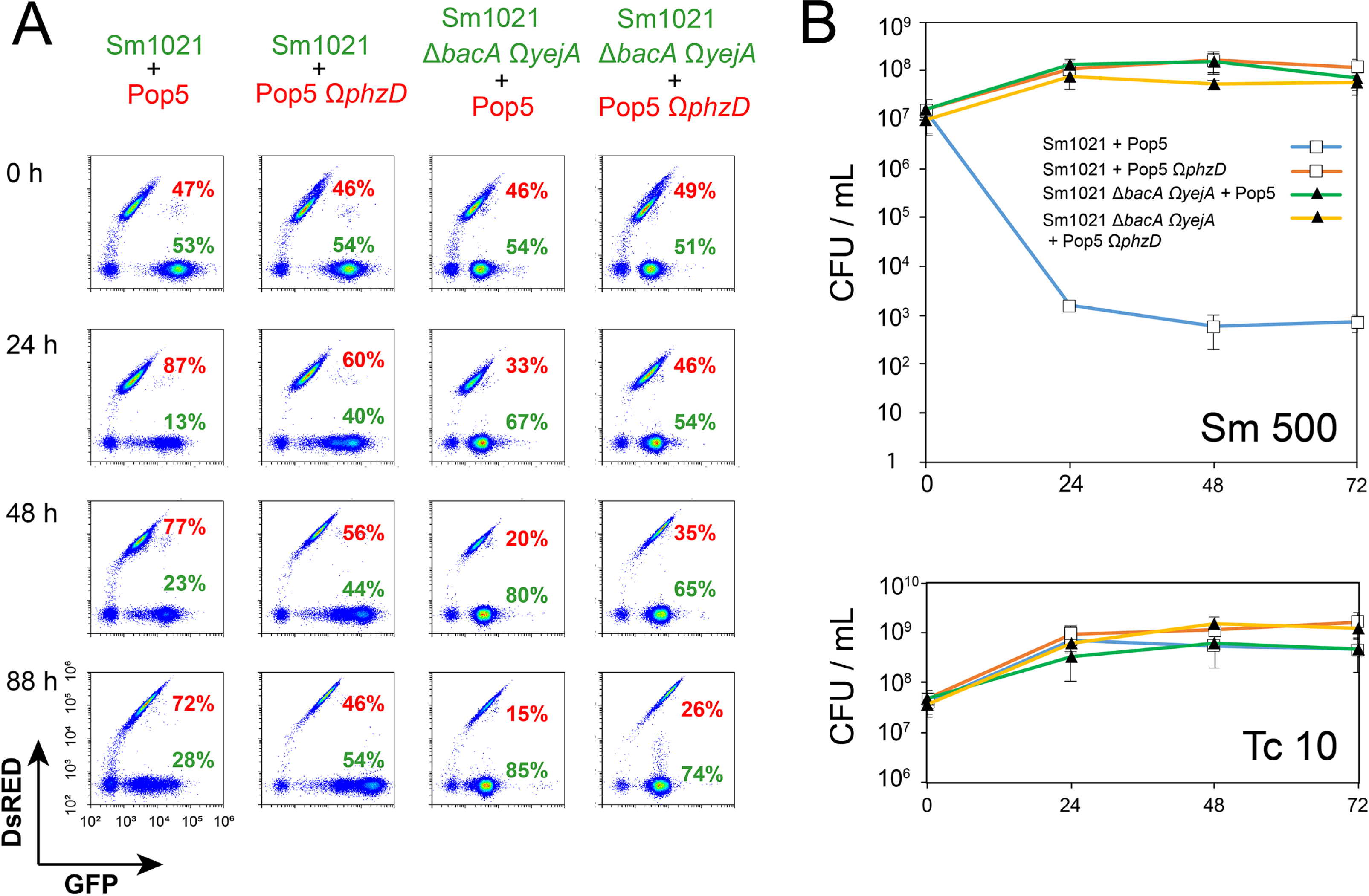
Sm 1021 lacking both YejABEF and BacA avoids the elimination by *Rhizobium* sp. Pop5 in co-cultivation experiments. **(A)** Identification by flow cytometry of Sm1021 derivatives and Pop5 derivatives in co-cultures grown during the indicated times. Dot plots show the GFP fluorescence in the x-axis and DsRed fluorescence in the y-axis for individual cells in the culture (dots). Note the higher GFP fluorescence of the Sm1021 strain carrying the pDG71 plasmid encoding GFP compared to the Sm1021 Δ*bacA* Ω*yejA* double mutant carrying a chromosome-inserted *gfp* copy. **(B)** CFU counts for the aliquots of two-strain mixtures sampled every 24 hours and spotted on Petri dishes with selective media (growth on Sm 500 reflects the number of *S. meliloti* Sm1021 CFUs, on Tc 10, the number of *Rhizobium* sp. Pop5 CFUs).

Though the flow cytometry analysis indicated a drop of the Sm1021 *wt* population in the mixture with Pop5 *wt*, the remaining fraction of PHZ-sensitive Sm1021 cells was relatively high (from 28 to 13%, depending on the time of cultivation). We decided to repeat the co-cultivation experiment using CFU counting on the selective media to assess the number of viable cells of each strain in mixed cultures. *Rhizobium* sp. Pop5 *wt* and *Rhizobium* sp. Pop5 Ω*phzD* transformed with the DsRed fluorescent protein-encoding plasmid pIN72 (Tc^R^) were mixed in a 1:1 ratio with either *S. meliloti* Sm1021 *wt* or Δ*bacA* Ω*yejA* (both Sm^R^) and cultivated for 4 days. In parallel, the growth of pure cultures of each strain was monitored. Culture aliquots were withdrawn at the start of the cultivation and every 24 hours afterward and used for CFU counting on tetracycline- or streptomycin-containing media (**Fig. 2B, Fig. S4B)**. After 24 h of co-cultivation of two *wt* stains, the number of Sm1021 *wt* viable cells dropped by almost four orders of magnitude. No increase in the number of *S. meliloti* CFUs was observed during further cultivation, while the number of *Rhizobium* sp. Pop5 CFUs increased by the end of the experiment. In contrast, for mixed cultures containing either transporter-deficient Sm1021 Δ*bacA* Ω*yejA* or PHZ-nonproducing *Rhizobium* sp. Pop5 Ω*phzD*, the CFU/mL numbers on both selective media increased ∼10-fold during the first 24 hours of cultivation and remained stable afterward. We conclude, that at our conditions, the transporter-deficient strain successfully avoided the action of PHZ produced by the *Rhizobium* sp. Pop5, which very efficiently eliminated the PHZ-susceptible *wt* Sm1021. The difference in total cell counts (by flow cytometry) and viable cell counts (CFU) suggests that initially Sm1021 *wt* cells multiply for several generations until the levels of PHZ in the co-culture are sufficiently high to kill the PHZ-susceptible bacteria and that dead bacteria persist for at least several days. The reduction of GFP-fluorescence we observed in these cells thus reflects their loss of viability.

### BacA and YejABEF homologs from other bacteria are capable of PHZ transport

Genes encoding the transporters homologous to BacA and YejABEF of *S. meliloti* (hereafter, referred to as BacA^Sm^ and YejABEF^Sm^) are widely distributed across the genomes of Alfa- and Gammaproteobacteria [17]. We constructed pSRK [23] shuttle vector-based plasmids harboring several such *bacA/yejABEF* homologs under the control of the inducible *lac* promoter. We selected transporters, which were previously shown to internalize peptide antibiotics (SbmA and YejABEF [17] of *E. coli*, NppA1A2BCD of *Pseudomonas aeruginosa* [24]) or to be required for the establishment of “host-symbiont” (BclA from *Bradyrhizobium* sp. [25]) and “host-pathogen” (BacA^Ba^ from *Brucella abortus* [26]) relationships (**Fig. 3A**). These constructs along with the empty pSRK vector control were transformed to the PHZ-resistant Sm1021 mut#1 described above. Colony formation on plates with and without PHZ in the presence of the inducer (1 mM IPTG) was monitored. With the exception of *yejABEF*^Ec^, strains expressing the genes of transporters were susceptible to PHZ, while the strain harboring the empty vector control was resistant, as expected (**Fig. 3B**). The PHZ-susceptible phenotype reversion observed for all but one strain indicates the ability of the tested transporters, with the exception of YejABEF^Ec^, to internalize PHZ.

**Figure 3.**
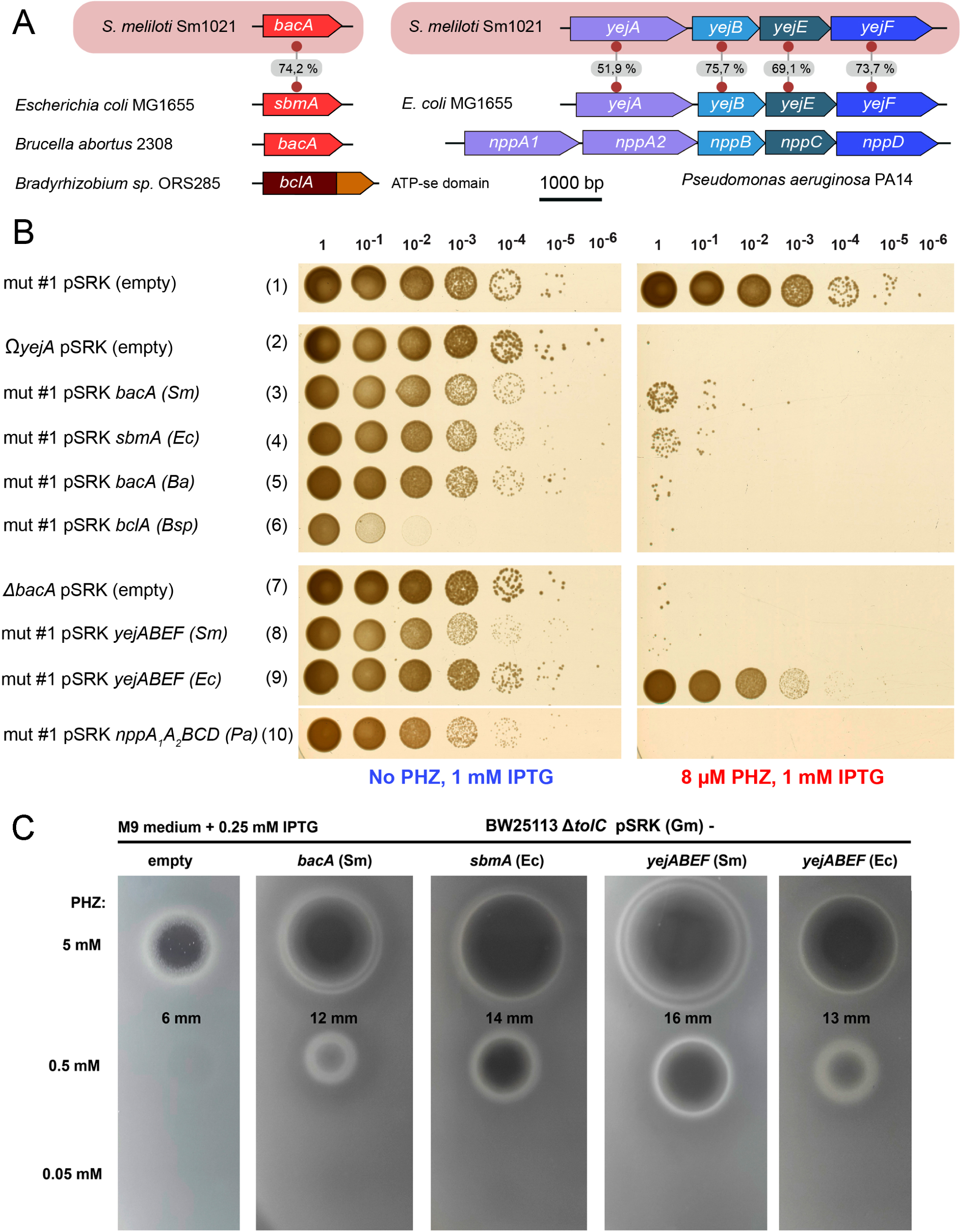
Resistance to PHZ can be reverted by the episomal expression of BacA, YejABEF or their orthologs from various Alfa- and Gammaproteobacteria. **(A)** Schematic representations of genes (*bacA*-like) and gene operons (*yejABEF*-like), which were chosen for the expression in PHZ-resistant Sm1021 (mut #1). Numbers on the grey background indicate the percentage of amino acid sequence similarity between the homologous proteins from *S. meliloti* Sm1021 and *E. coli* MG1655. **(B)** The results of the CFU assay on the Petri dishes with 8 μM PHZ and without PHZ for the strains of Sm1021 expressing different genes (operons) of transport proteins upon IPTG induction. Note that the CFU count for the *yejABEF(Ec)*-expressing strain (lane 9) is approximately 10 times lower in the presence of PHZ compared to the medium without PHZ and this is not observed for the empty plasmid control (lane 1). **(C)** The inhibition zones from 5 μL of PHZ spotted in 3 different concentrations (5, 0.5, and 0.05 mM) on top of a loan of *E. coli* BW25113 Δ*tolC* cells expressing the genes of different transport proteins or containing empty pSRK vector. The sizes of zones for 5 mM PHZ are indicated. IPTG, isopropyl β-D-1-thiogalactopyranoside. Note that the use of high PHZ concentrations was required because of the intrinsic high resistance of *E. coli* to PHZ.

The fact that expression of YejABEF^Ec^, a close homolog of YejABEF^Sm^, led to only very moderate increase in PHZ-sensitivity (**Fig. 3B line (9)**), may be due to the lower affinity for PHZ of the periplasmic ABC-transporter subunit YejA^Ec^ or because of poor assembly of the multisubunit transporter in a heterologous Sm1021 host. To distinguish between these possibilities we wanted to test the ability of YejABEF^Ec^ to transport PHZ in its native host. However, *E. coli* is naturally resistant to PHZ in concentrations 100 times higher than those inhibitory for rhizobia [5]. A possible contributing factor to PHZ-resistance of *E. coli* is TolC, which is a major outer membrane multidrug efflux protein that can expel various compounds from the cell [27]. Indeed, an *E. coli tolC* deletion mutant has an increased sensitivity to PHZ [5]. Similarly, the PHZ MIC for a Sm1021 Ω*tolC* mutant is at least eight times lower than for the *wt* (**Table 1**). To test the ability of *E. coli* peptide transporters to import PHZ in their native host *in vivo*, we transformed *E. coli* BW25113 Δ*tolC* with pSRK-based *bacA*^Sm^, *sbmA*^Ec^, *yejABEF*^Sm^, or *yejABEF*^Ec^ expression plasmids. Lawns of resulting strains and the empty plasmid-bearing control strain grown in the presence of IPTG were spotted with drops of PHZ solutions and the appearance of growth inhibition zones was monitored. Interestingly, the expression of any of the four transporters either from Sm1021 or *E. coli* led to an increase in the inhibition zone sizes compared to control (**Fig. 3C**). Thus, the lack of PHZ sensitivity upon expression of *yejABEF*^Ec^ in Sm1021 is more likely caused by protein misfolding or inefficient expression.

### BacA and YejABEF are involved in the internalization of the PHZ-unrelated thiazole-containing antibiotic bleomycin

Next, we aimed to determine whether compounds other than PHZ are also internalized via the same pair of *S. meliloti* transporters. Previously, *in vivo* experiments with the *bacA* null mutant demonstrated that BacA^Sm^ contributes to the sensitivity of *S. meliloti* to the thiazole-containing peptide-polyketide hybrid antibiotic bleomycin A2 (BLM) [11]. While the direct uptake of BLM by BacA^Sm^ has been reported previously [10], the observed partial resistance to BLM of the *bacA* mutant pointed towards the involvement of an additional BacA-independent pathway for BLM internalization [11]. We determined the MICs of BLM against *wt* Sm1021 along with single and double *bacA* and *yejABEF* mutants using a broth microdilution assay. In agreement with published data, compared to the *wt*, the Δ*bacA* Sm1021 mutant was twice less susceptible to BLM. Both Ω*yejA* and Ω*yejE* single mutants had similarly increased resistance to BLM (a MIC of 0.66 μM compared to 0.33 μM for the *wt*, see **Table 1**). We, therefore, conclude that the YejABEF provides an alternative pathway for BLM uptake. Remarkably, the double mutants Δ*bacA* Ω*yejA* and Δ*bacA* Ω*yejE* were 16 times more resistant to BLM than the *wt* (**Table 1**). This data indicate that YejABEF- and BacA-mediated pathways of BLM uptake are independent of each other. While these two transporters are largely responsible for the BLM sensitivity of Sm1021, the inhibition of double mutants growth seen at high concentrations of BLM may be due to the function of yet another low-affinity transport system that remains to be identified or to a low-efficient transporter-independent uptake mechanism.

### In vitro transport assay shows the internalization of PHZ via BacA and SbmA

The expression of BacA^Sm^ and all its tested homologs (SbmA, BacA^Ba^, BclA) reverted the PHZ-susceptible phenotype in PHZ-resistant double mutants, which is consistent with the promiscuous peptide uptake by these transporters [10–14,25,28]. While there is no *in vitro* system to study transport by YejABEF, we have previously developed a system to monitor substrate transport by SLiPTs including BacA^Sm^ and SbmA^Ec^ [10]. We used this liposome transport assay to provide direct evidence of the ability of BacA^Sm^ to internalize PHZ. The assay monitors pyranine fluorescence inside the liposomes, which decreases upon acidification. We have previously shown that peptide transport by BacA^Sm^ is coupled to the proton gradient [10] and decrease in fluorescence of liposomes containing BacA^Sm^ in the presence of PHZ indicates that the peptide can be transported into the liposomes (**Fig. 4A**). Our data show that SbmA, the *E. coli* ortholog of BacA, is also capable of transporting PHZ (**Fig. 4B**). Noteworthy, BacA and SbmA show different transport kinetics; BacA appears to have an initial delayed uptake that could be either due to the PHZ recognition within the cavity or conformational changes associated during the transport cycle. Be that as it may, the result is consistent with the interchangeable role of SbmA and BacA proteins (**Fig. 3B**).

**Figure 4.**
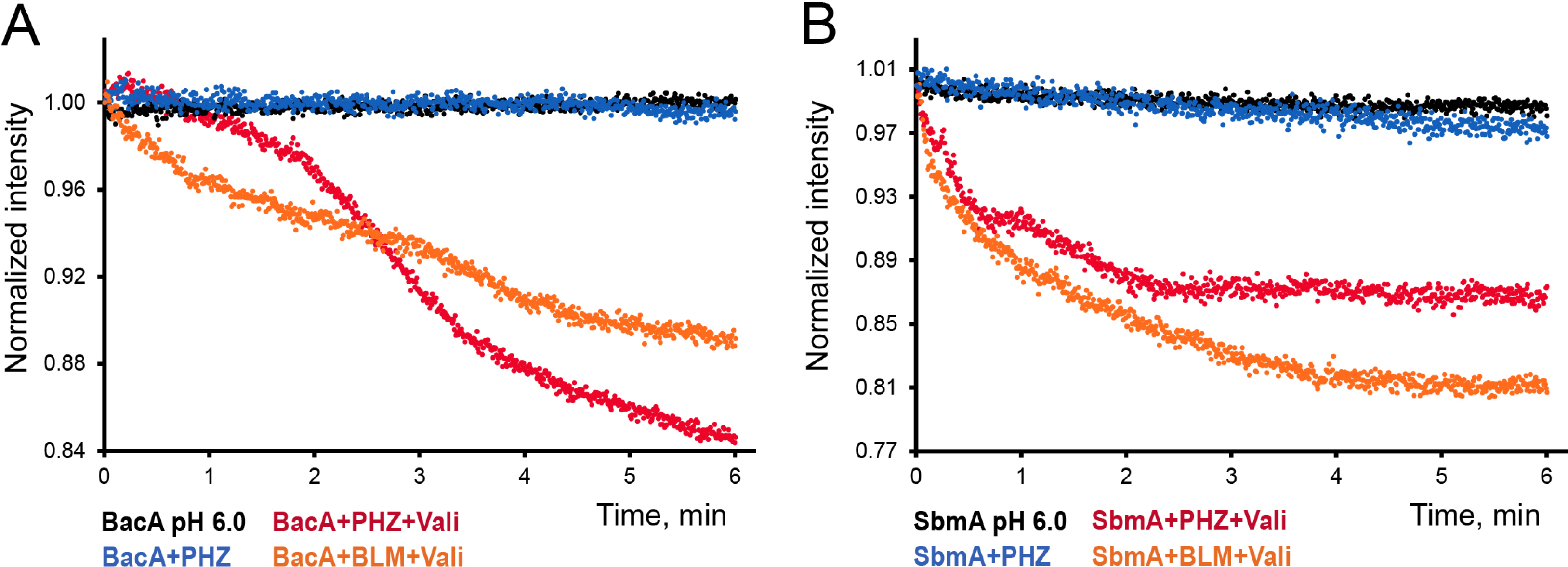
Results of the *in vitro* liposome transport assay, showing the BacA-mediated. **(A)** and SbmA-mediated **(B)** import of PHZ. Bleomycin (BLM) was used as a positive control. Vali, valinomycin. For the description of the assay, see the Methods section.

### The structure of YejA^Sm^ and YejA^Ec^

To get insights into the recognition mechanism of PHZ by the YejABEF transporter, we determined the structures of the substrate-binding proteins (SBPs) YejA^Sm^ and YejA^Ec^. For other ABC transporters involved in peptide uptake, it has been shown that SBPs determine the specificity of transport [29–31]. SBPs exist in either open (apo) or closed (ligand-bound) conformations [32]. While we purified and crystallized each protein in the absence of externally added ligands, in both cases short peptides were found to be associated with the protein. Our numerous attempts to obtain either YejA free of bound peptides, including denaturation/renaturation of purified proteins and/or expression in the periplasm, were unsuccessful.

The structures of YejA^Sm^ and YejA^Ec^ proteins with bound peptides were determined at 1.58 Å and 1.65 Å resolution, respectively. The numbering used below for the description of residues corresponds to mature YejA^Sm^ and YejA^Ec^ lacking their signal peptides; *i*.*e*. residue number 1 in YejA^Sm^ and YejA^Ec^ is encoded by codons of Glu31 and Ala21 in *yejA*^*Sm*^ and *yejA*^*Ec*^, respectively. Both YejA^Sm^ and YejA^Ec^ share a highly similar fold consisting of two lobes, each formed by a central β-sheet flanked by α-helices (**Fig. 5A** and **5B**): they display a root-mean-square deviation (rmsd) of 1.62 Å for 543 Cα atoms (**Fig. 5C**). The bigger lobe (lobe 1) consists of residues 1-290 and 558-589 (1-279 and 545-584) and the smaller one (lobe 2) comprises residues 301-552 (284-538) for YejA^Sm^ (YejA^Ec^). Two short segments (residues 291-300 and 553-557 in YejA^Sm^) serve as hinges connecting the two lobes (**Fig. 5A** and **5B**, black).

**Figure 5.**
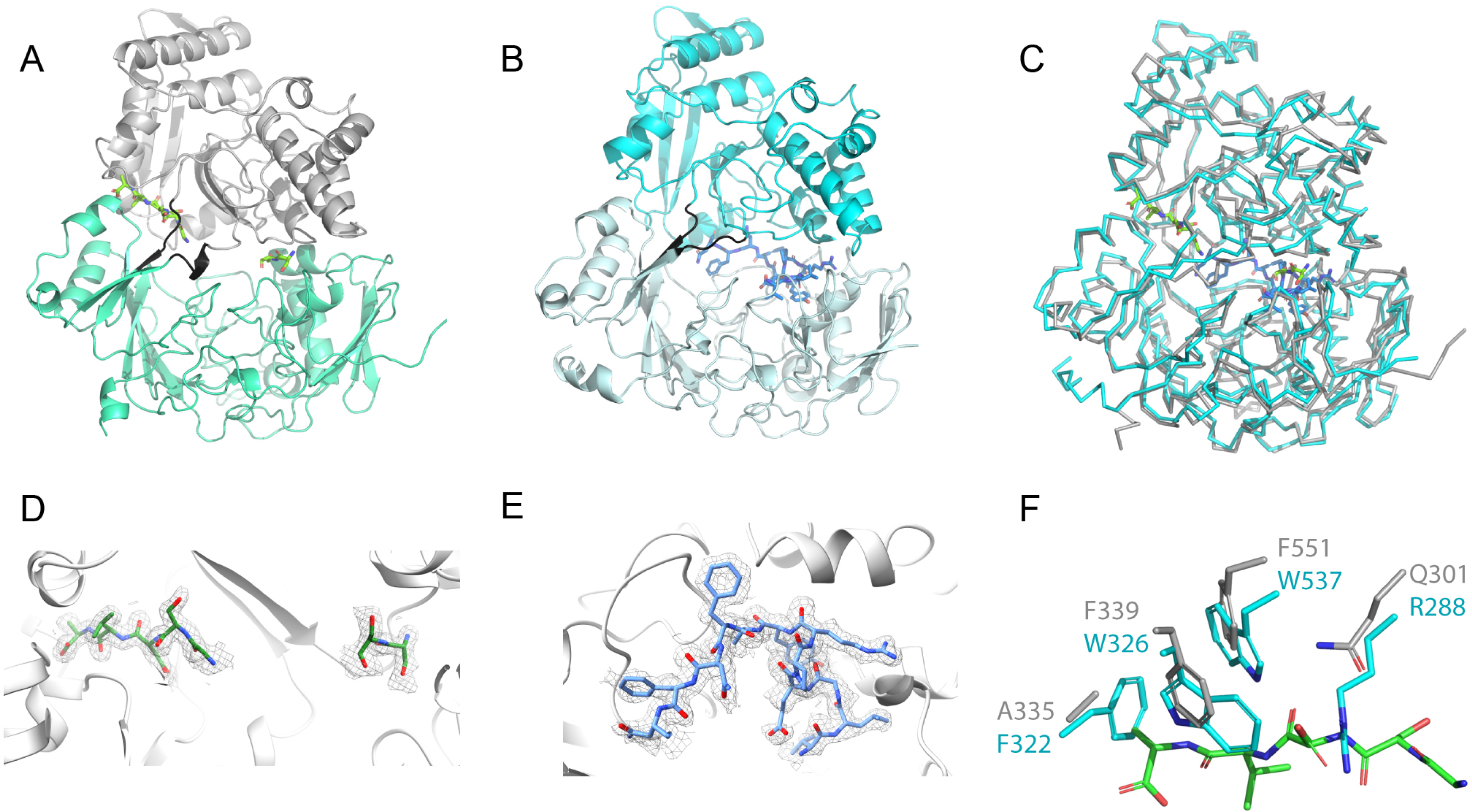
Crystal structures of YejA^Sm^ and YejA^Ec^ with fortuitous peptides bound. **(A)** Ribbon representation of YejA^Sm^ (PDB ID: 7Z8E) with two distinct degraded peptides (a pentapeptide and a dipeptide in stick format) shown in green, bound at the interface between the two lobes (grey and blue). The short hinge region is shown in black. (**B**) YejA^Ec^ (PDB ID: 7Z6F) with a dodecapeptide shown in blue in stick format, bound at the interface between the two lobes (cyan and grey). The short hinge region is shown in black. (**C**) Structural superimposition of YejA^Sm^ (grey) and YejA^Ec^ (cyan) structures. (**D**) Electron density for the bound degraded peptides in the YejA^Sm^ binding pocket. (**E**) Electron density for the bound dodecapeptide in the YejA^Ec^ binding pocket. The 2Fo-Fc electron density map for both proteins is contoured at 1*σ* and is shown as grey mesh. (**F**) Structural comparison between YejA^Sm^ (grey) and YejA^Ec^ (cyan) around the pentapeptide (green) bound to YejA^Sm^. Side chains of the residues are labelled and shown as sticks.

YejA possesses a typical cluster C fold within the SBP structural classification [32], as reported by SSM-EBI (http://www.ebi.ac.uk/msd-srv/ssm). Indeed, obtained YejA structures resemble those of oligopeptide-binding SBPs such as the Cu(I)-methanobactin complex-binding MbnE from *Methylocystis parvus* (PDB ID: 5ICQ) [33], the oligopeptide-binding AppA from *Bacillus subtilis* (PDB ID: 1XOC) [34], and the oligopeptide-binding OppA from *Lactococcus lactis* (PDB ID: 3DRF) [30]. Noteworthy, although all these SBPs share a similar fold, their oligopeptide-binding sites are distinct. Both YejA structures adopt a closed conformation due to the binding of fortuitous oligopeptides at the interface between the two lobes.

The bound oligopeptides could originate from YejA degradation and/or peptides from *E. coli* during overexpression. Nonetheless, the determined YejA crystal structures correspond to the full length proteins without signs of degradation. A dodecapeptide (VLGEPRYAFNFN) was built in YejA^Ec^, guided by the electron density (**Fig. 5D**), the assignment of the peptide side chains bound to YejA^Sm^ was more challenging (**Fig. 5E**). A dipeptide (SS) and a pentapeptide (GSDVA) were built, each at different places in the closed interface. Electron density linking the two peptides was missing emphasizing that a population of different peptides might have bound to the peptide-binding site. Consequently, the resultant crystal structure of YejA^Sm^ is an average of different peptides in the crystal lattice. The pentapeptide observed in YejA^Sm^ binds in a pocket that is larger than in YejA^Ec^ where bulkier residues of YejA^Ec^ (Arg288, Phe322, Trp326, Trp537 versus Gln301, Ala335, Phe339, Phe551 in YejA^Sm^) would prevent peptide presence in this interface area (**Fig. 5F**). In contrast, the dipeptide bound to YejA^Sm^ is located near the beginning of the dodecapeptide bound to YejA^Ec^ (**Fig. 5C**). Binding of the pentapeptide to YejA^Sm^ is almost exclusively due to hydrogen bonds formation between the peptide backbone and the protein (10 interactions out of 12) and that of the dipeptide is only via its backbone, meaning that the binding is unspecific. In YejA^Ec^, the dodecapeptide makes half (9 out of 18) of its interactions with the protein through the backbone. Eleven side chains of YejA^Ec^ indicated by red triangles in the structure-based sequence alignment (**Fig. S5**) are involved in the dodecapeptide binding. Only two of them are conserved in YejA^Sm^ suggesting that the dodecapeptide would not be accommodated in YejA^Sm^.

## DISCUSSION

Here, we demonstrate that PHZ utilizes a unique dual transporter uptake mechanism to cross the inner membrane of *S. meliloti* Sm1021 via two unrelated peptide transporters, BacA and YejABEF. BacA adopts the SLiPT fold and is powered by the proton-motive force [10], while YejABEF is an ABC transporter, powered by ATP hydrolysis. BacA of *S. meliloti* was previously shown to internalize the thiazole-containing antibiotic BLM [11], while SbmA, its ortholog in *E. coli*, is required for the uptake of azole-modified RiPPs microcin B17 [13] and klebsazolicin [16]. Thus, PHZ provides another example of an azole-containing molecule internalized through a SLiPT. The exact mechanism of peptide recognition by SLiPTs is still unknown. Available SLiPT structures reveal a large cavity where structurally unrelated antimicrobial peptides are thought to bind [10]. The large size of the likely ligand-binding site explains the observed promiscuity of SLiPTs, which in addition to azole-containing peptides also transport lasso-peptides [35], unmodified proline-rich peptides [15], NCR plant peptides (see below) [25], and a number of other natural [36] and artificially designed substrates [37,38]. *E. coli* YejABEF is the only entry point for the heptapeptide-nucleotide antibiotic microcin C [17]. Interestingly, when the peptide moiety of microcin C was artificially increased to 25 amino acids, import through SbmA became also possible and both systems contributed to the compound internalization [39]. These data are consistent with our observations on the uptake of PHZ in Sm1021, whose length (27 amino acids) is comparable to that of the 25-amino acid long derivative of microcin C.

In *S. meliloti*, BacA and YejABEF are involved in the transport of nodule-specific cysteine-rich peptides (NCRs) [19,25]. NCRs are a diverse family of ribosomally-synthesized defensin-like peptides encoded in the genomes of some plants (such as *Medicago*) with lengths varying usually from 24 to 65 amino acids [40,41]. Cationic NCRs exhibit antimicrobial activity by targeting cell membranes of both Gram-positive and Gram-negative bacteria [42]. A mixture of NCRs secreted by plant cells in the nodules allows the host to control the development and the composition of the endosymbiotic rhizobial population located within the nodule plant cells [40,43]. Since internalization and subsequent intracellular degradation of NCRs protects the bacteria, transporters involved in NCR uptake play a key role in establishing functional symbiosis. An *S. meliloti* mutant lacking BacA is unable to form functional nodules on the roots of the legume *Medicago truncatula* and dies rapidly in the plant nodule cells due to the inhibitory action of NCRs [44]. Likewise, *S. meliloti* strains with the deletions of any one of the *yejABEF* operon genes form highly abnormal hypertrophied cells inside the nodules due to increased susceptibility to NCRs [19]. Interestingly, the *yej* genes in pathogenic *Salmonella* and *Brucella* contribute to virulence by protecting bacteria from host-produced membrane-targeting antimicrobial peptides, most likely via their internalization [45,46]. Note that in the case of membrane-attacking peptides such as the NCRs or the immunity antimicrobial peptides of animals, the BacA and YejABEF transporters provide protection contrary to the case of peptides with intracellular targets such as PHZ or BLM, for which the activity of these transporters renders bacteria sensitive.

PHZ is an exciting example of an NCR-unrelated antirhizobial compound internalized via two independent pathways. The dual-entry mode of PHZ by itself dramatically decreases the rate of resistance compared to compounds with a single entry point. Moreover, bacteria that managed to acquire PHZ resistance through mutations in both transporters will be unable to develop functional nodules, for which the uptake of membrane-targeting NCRs is essential. Since passage through symbiosis and massive multiplication inside the legume nodules, followed by the return of bacteria into the soil at the end of the nodules’ lifetime, is a key mechanism of rhizobial spread in the environment, the loss of any one of these transporters will be evolutionary disfavoured. As such, PHZ may have potential as a biocontrol agent for agriculture, for example, in legume crop inoculation strategies with elite rhizobium strains.

The obtained structures of YejA^Sm^ and YejA^Ec^ showed that these proteins are oligopeptide-binding SBPs that can interact with fortuitous peptides in a non-specific manner. Their natural ligands still remain unknown, with microcin C being the only known biologically relevant ligand of YejABEF^Ec^. Our work shows that overexpression of the YejABEF^Ec^ transporter increases the sensitivity of *E. coli* to PHZ, which makes it another substrate. In our structure, YejA^Sm^ appears to bind multiple peptides, which may be derived from the proteins of the heterologous expression host, *E. coli*. The recognition of multiple degradation products has been reported for peptide transporters AppA from *B. subtilis* [34], and OppA from *L. lactis* [30], though these transporters have not been linked with antimicrobial peptide recognition. Whatever the role of the peptides bound to YejA^Sm^ and YejA^Ec^, their presence precluded us from solving the structures with either PHZ or microcin C. Presumably, the closed conformation of peptide-bound SBPs and the tight binding of the peptides that have been selected from the pool available in the cytoplasm did not allow for the exchange of bound peptides with externally added antibiotics.

PHZ displays low-micromolar MICs against rhizobia closely-related to the producing strain *Rhizobium* sp. Pop5, while multiple strains of rhizobia (for instance, those of the genus *Mesorhizobium*) along with various Gammaproteobacteria are resistant to even high concentrations of PHZ [5]. Here, we demonstrate that orthologs of both BacA and YejABEF are capable of reverting the PHZ-resistant phenotype once expressed heterologously, which implies that they can perform PHZ internalization in their native hosts as well. In fact, we show that *E. coli* SbmA and YejABEF can internalize PHZ. Since previously we demonstrated that *E. coli* ribosomes are highly sensitive to PHZ *in vitro* [5], it remains unclear, why the PHZ MIC for *E. coli* is at least two orders of magnitude higher compared to that for *S. meliloti*. We propose that there may be additional factors contributing to the lower sensitivity of *E. coli*, among which there is the active TolC-mediated export of PHZ [5]. However, this can not be the sole factor since here we show that a *tolC* mutant of *S. meliloti* has a nanomolar MIC for PHZ, which is at least a hundred-fold lower than that of the *E. coli* Δ*tolC* mutant. These observations demonstrate that the sensitivity of a certain strain to a given antimicrobial is a complex function, which cannot be simply inferred based on our knowledge of the target structure and the presence of certain import and export systems. Additional factors such as the level of expression of genes coding for import and export machinery confounded by minor differences in specificities/kinetics revealed in our *in vitro* liposome uptake experiments may result in dramatic differences in the final MIC values observed.

It is not yet known whether PHZ provides a competitive advantage to its producer either in soil or in the competition for root nodulation, as it was previously shown for another antirhizobial RiPP trifolitoxin [8,9]. However, our co-cultivation studies show that this is clearly the case in laboratory conditions. We identified BGCs virtually identical to the *Rhizobium* sp. Pop5 PHZ BGC in multiple recently sequenced genomes of Alphaproteobacteria and one genome of a Betaproteobacterium sampled around the globe (**Fig. S6A**). The occurrence of clusters guiding the biosynthesis of close PHZ homologs (**Fig. S6B, C**) across bacteria of multiple genera (including both nodule-forming and plant-associated rhizosphere species, **Table S4**) derived from geographically distant sampling sites may indicate that the ability to produce PHZ-like compounds may be a common and efficient mechanism that provides bacteria with competitive advantages in complex environmental niches.

## METHODS

### Bacterial strains and growth conditions

Bacterial strains used in the study are listed in **Table S1**. For *S. meliloti*, YEB medium (per 1L: 5 g peptone, 5 g beef extract, 5 g sucrose, 1 g yeast extract, 0.4 g MgSO_4_·7H_2_O, pH 7.5) was used if not specified differently. *Rhizobium* sp. Pop5 was cultivated in YM medium (per 1 L: 10 g mannitol, 0.5 g K_2_HPO_4_, 0.2 g MgSO_4_, 0.1 g NaCl, 1 g yeast extract, pH 6.8). *E. coli* strains were grown in LB medium (per 1 L: 5 g NaCl, 10 g tryptone, and 5 g yeast extract) or 2xYT medium (per 1 L: 5 g NaCl, 16 g tryptone, and 10 g yeast extract). Rhizobia were cultivated at 28°C, *E. coli* strains at 37°C. Antibiotics were used in the following final concentrations: ampicillin (Ap), 100 μg·mL^-1^; kanamycin (Km), 50 μg·mL^-1^ for *E. coli* and 100 μg·mL^-1^ for rhizobia; tetracyclin (Tc), 10 μg·mL^-1^; chloramphenicol (Cm), 34 μg·mL^-1^; spectinomycin (Sp), 25 μg·mL^-1^; streptomycin (Sm), 500 μg·mL^-1^; and gentamycin (Gm), 50 μg·mL^-1^.

### Sm1021 transposon library construction

The strain *S. meliloti* Sm1021 carrying resistance to Sm was used for transposon mutagenesis and was cultured in YEB medium supplemented with Sm at 28°C. The *E. coli* MFD*pir* strain [47] (Δ*dapA*-derivative, auxotroph for diaminopimelic acid (DAP) synthesis) carrying the plasmid pSAM_Ec [48] was used as a donor strain for the transposon mutagenesis, and cultured in LB supplemented with 300 μg·mL^-1^ of DAP and Km at 37°C. The donor strain *E. coli* MFD*pir* pSAM_Ec and the recipient strain *S. meliloti* Sm1021 were grown in 50 mL cultures at 180 rpm until the exponential growth phase at a final OD_600nm_ of 1. The cultures were washed twice (centrifugation at 4000 rpm for 10 minutes at room temperature) with fresh medium without antibiotics. The pellets were resuspended in fresh medium without antibiotics to obtain a final OD_600nm_ of 50. For conjugation, the donor strain and the recipient strain were mixed at a ratio 1:1. Multiple 100 μL drops of the bacterial mix were spotted on YEB agar plates supplemented with 300 μg·mL^-1^ of DAP and incubated at 28°C. After 6 hours of incubation, allowing conjugation of the pSAM_Ec plasmid from the donor *E. coli* strain to the Sm1021 recipient strain and transposition of the transposon into the genome of the target strain, the spots were resuspended in YEB medium, a dilution series was plated on selective medium carrying Sm and Km and subjected to CFU counting to assess the number of individual bacterial mutants obtained by the mutagenesis. In parallel, the remaining bacterial suspension was spread on YEB agar plates supplemented with Sm and Km to obtain the *S. meliloti* Sm1021 transposon mutant population. After 2 days of incubation at 28°C, the transposon library was resuspended from the agar plates in fresh liquid YEB medium. The suspension was adjusted to 20% glycerol, aliquoted and stored at -80°C.

### Selection of phazolicin-resistant mutants

100 μL of the *S. meliloti* Sm1021 Tn-library prepared as described above with the cell concentration of approximately 1*10^8^ cells·mL^-1^ were plated on two Petri dishes with YEB medium containing Sm, Km and 20 μM of PHZ (∼20xMIC). Petri dishes were incubated for 48 hours at 28°C. Obtained colonies were restreaked on a Petri dish with fresh PHZ-containing medium to confirm the resistance phenotype.

### Whole-genome sequencing, identification of transposon insertion positions

DNA was extracted from 3 mL overnight cultures of *S. meliloti* Sm1021 resistant mutants using GeneJET Genomic DNA purification kit (ThermoFisher) according to the manufacturer protocol. NGS libraries were prepared using NEBNext Ultra II DNA Library Prep kit (NEB). DNA sequencing was performed on Illumina MiSeq with the 250+250 bp paired-end protocol. Library preparation and sequencing were performed at Skoltech Sequencing Core Facilities. Raw reads were filtered and trimmed with Trimmomatic [49], genome assembly was performed with SPAdes [50]. Identification of transposon insertion positions was performed with a stand-alone BLAST using the Km^R^ gene sequence as a bait [51]. NC_003047 genome annotation was used as a reference.

### Construction of PHZ-nonproducing strain Rhizobium sp. Pop5 ΩphzD

To obtain a *Rhizobium* sp. Pop5 mutant with the disruption of the *phzD* gene (YcaO domain-containing cyclodehydratase, locus tag: RCCGEPOP_21747, protein GenBank accession number: EJZ19165.1), a 566 bp internal fragment of the gene was PCR amplified and cloned into the plasmid pVO155nptIIgfp (pVO155 plasmid [52] derivate with constitutively expressed *gfp* gene; does not replicate in *Rhizobium* spp.) between *SalI* and *XbaI* restriction sites. The resulting construct was introduced into *Rhizobium* sp. Pop5 via triparental mating with the helper strain HB101 pRK600 [53]. The cells with the plasmid integrated into the genome were selected on YM medium with Km. As there is no resistance marker in the genome of *Rhizobium* sp. Pop5, we did not perform a counter-selection of the *E. coli* donor, which could be easily distinguished from *Rhizobium* based on the morphology of the colonies growing on the solid YM medium.

### Construction of Sm1021 double mutants

Generalized transduction by *S. meliloti* Sm1021 phage ϕM12 was used to obtain the double mutants lacking both functional BacA and YejABEF importers. *S. meliloti* Sm1021 Δ*bacA* served as a donor, while Sm1021 Ω*yejA* and Sm1021 Ω*yejE* were used as recipient strains. The procedure was performed as described earlier [54]. Briefly, 5 mL of Sm1021 Δ*bacA* donor strain culture grown overnight at 30°C in LB/MC medium (LB medium + 2.5 mM CaCl_2_+2.5 mM MgSO_4_) supplemented with Sp, was inoculated by the phage in cell:phage ratio 1:1. The mixture was incubated overnight with shaking at 30°C, then sterilized by the addition of 150 μL of chloroform and cleared from the remaining cell debris by centrifugation (10,000 rpm, 10 min). The obtained lysate was then used to inoculate 1 mL of the overnight culture of the recipient strains grown in LB/MC up to cell:phage ratio of 2:1. The obtained mixtures were incubated for 30 min at room temperature and pelleted (8,000 rpm, 2 min). The pellet was washed with 1 ml of TY medium (per 1L: tryptone 5 g, yeast extract 3 g) and resuspended in fresh TY. The suspensions were plated on the TY agar plates supplemented with Sp and Km to select for transductants carring both resistance markers in the genome. A mixture lacking the lysate served as a negative control. The colonies obtained were screened using PCR with primers specific to *bacA* and *yejA* or *yejE* to confirm the genotype.

### Molecular cloning procedures

**Table S1** includes the list of plasmids used in the study. Oligonucleotide primers are listed in **Table S2**. Molecular cloning of the genes encoding BacA-related and YejABEF-related transporters into the pSRK vector [23] was performed either by conventional restriction enzyme digestion and ligation protocol (restriction sites are specified for each gene in the corresponding primer names) or by Gibson Assembly protocol (NEB). For Gibson Assembly pSRK plasmid was PCR-amplified with primers pSRK_GA_F and pSRK_GA_R and treated with DpnI restriction endonuclease (ThermoFisher).

### Co-cultivation competition experiments and flow cytometry

For flow cytometry analysis of competition experiments, the GFP-expressing strains Sm1021 pDG71 and Sm1021 Δ*bacA* Ω*yejA* and the DsRed-expressing strains Pop5 pIN72 and Pop5 Ω*phzD* pIN72 were used. Pre-cultures of the strains, grown in YM medium with appropriate antibiotics, were washed and diluted in fresh YM medium without antibiotics to OD_600nm_=0.1. Single strain cultures or 50%:50% mixtures were prepared from these fresh suspensions to reach a final OD_600nm_=0.4 in YM medium without antibiotics. Aliquots were taken from these cultures at t=0 h, t=24 h, t=48 h, and t=88 h for analysis by flow cytometry.

Flow cytometry was performed with a CytoFLEX (Beckman Coulter) instrument. Gating on the bacterial particles was done using forward and side scatters, fluorescence levels of the bacterial particles were acquired using the preset GFP and DsRed channels of the instrument. For each measurement, 50000 events were recorded and plotted in the dot plots of **Fig. 2A** and **Fig. S4A**. Data analysis and representation in dot plots were performed with the CytExpert version 2.4.0.28 software (Beckman Coulter).

The preparation of mixtures for the co-cultivation experiment with CFU number monitoring was performed essentially as described above. Sm1021 *wt* was used instead of Sm1021 pDG71 to eliminate the effect of the plasmid maintenance on the growth of the culture. Aliquotes were taken from the cultures at t=0 h, t=24 h, t=48 h, and t=72 h. Ten-fold dilution series were prepared from these aliquotes, 5 μL of each dilution was spotted on YM plates without antibiotics or with either Sm or Tc. CFU counting was performed after 48 hours of incubation at 28°C.

### Broth microdilution assays and MIC determination

Precultures of the *wt S. meliloti* Sm1021 and mutants were grown in YEB medium with Sm. Overnight grown cultures were diluted to OD_600nm_ of 0.2 in fresh YEB medium with Sm and grown until OD_600nm_ of 1. The cells were pelleted by centrifugation and resuspended in YEB medium without antibiotics until OD_600nm_ of 0.05. The cells were dispatched by 150 μL in a 96-well plate, except for the first column, which contained 300 μL of cultures. PHZ was added to the first column to a final concentration of 20 μM or BLM to a final concentration of 20 μg·mL^-1^. Two-fold serial dilutions in the subsequent columns were obtained by serial transfer of 150 μL to the next column and mixing by pipetting up and down. No peptide was added to the last column of the 96-well plate. The 96-well plates were incubated in a SPECTROstar Nano plate incubator (BMG LABTECH). The growth of the cultures in the wells was monitored by measuring the OD_600nm_ and data points were collected every hour for 48 hours. Plates were incubated at 28°C with double orbital shaking at 200 rpm. Data and growth curves were analyzed using Microsoft Excel. The assay was performed in triplicate for both PHZ and BLM.

### Colony-forming unit (CFU) assay

CFU counting was used to access the sensitivity of strains to the action of PHZ as a complementary method to the broth microdilution assay, as it allows to identify the occurrence of resistant clones, which appear as colonies growing in the undiluted to hundred-fold diluted samples. For CFU counting the overnight cultures of selected Sm1021 derivatives were diluted with fresh YEB medium with relevant antibiotics added to the OD_600nm_=0.2 and allowed to grow to the OD_600nm_=0.6 at 28°C with shaking. Then the cultures were adjusted to OD_600nm_=0.2 and ten-fold dilution series of the obtained cell suspensions were prepared, 5 μL of each dilution was spotted on YEB plates supplemented with either Sm (negative control) or Sm and 8 μM of PHZ. Each experiment was performed in triplicate using independent starting cultures inoculated with single colonies of the corresponding strains. For the strains carrying the pSRK plasmids with a panel of *bacA*/*yejABEF* orthologs under the control of the *lac* promotor, 1 mM IPTG was added to the YEB medium to induce the expression. Plates were incubated for 3 days at 28°C, after which the number of CFUs was counted.

### Production and purification of phazolicin

Phazolicin was purified from the cultivation medium of *Rhizobium* sp. Pop5 following the protocol described previously [5]. Briefly, the procedure included solid-phase extraction on an Agilent HF Bond Elut LRC-C18 Cartridge, followed by reverse-phase HPLC purification on a Luna PREP C18 column.

### In vitro transport assays with BacA and SbmA-containing liposomes

SbmA and BacA proteins for functional assays were purified and reconstituted in liposomes using the rapid dilution method as previously described [10]. For the transport assay, 15 μM (final concentration) of liposomes were placed in a cuvette and quickly mixed with 1.0 mL of outside buffer (5 mM HEPES pH 6.8, 1.2 mM KCl, and 2 mM MgSO_4_) containing BLM (100 μM) or PHZ (50 μM) with or without valinomycin (1 μM). Data were recorded using a Cary Eclipse Fluorescence Spectrophotometer (Agilent Technologies) at the following setting: 460 nm excitation, 510 nm emission, 5 nm slit width, and 0.5 sec resolution for 6 min.

### Cloning, expression and purification of YejA^Sm^

YejA^Sm^ signal peptide prediction was performed using SignalP 5.0 [55]. *yejA*^Sm^ lacking the fragment encoding the first 30 amino acids (signal peptide) was PCR-amplified from Sm1021 genomic DNA and cloned into a pET29b(+) vector (Novagen) generating a C-terminal 6His-tag. The resulting plasmid pET29-*yejA*^*Sm*^-CHis6 was electroporated into *E. coli* Rosetta 2 (DE3) pLysS. Two liters of 2xYT medium supplemented with Km and Cm were inoculated with 20 mL of an overnight culture of the obtained strain. Cells were grown at 37°C, 180 rpm to an OD_600nm_=0.7, then induced with 0.5 mM IPTG and incubated at 28°C for another 5 hours. The cultures were cooled down on ice, pelleted (4000 rpm, 20 min, 4°C), and frozen in liquid nitrogen.

Cells were resuspended in 80 mL of Lysis Buffer (50 mM Tris-HCl pH 8.0, 300 mM NaCl, 10% glycerol, 20 mM imidazole) supplemented with homemade purified DNAse and protease inhibitors cocktail (Sigma-Aldrich) and disrupted by sonication. After centrifugation (17000 rpm, 25 min, 4°C), the supernatant was loaded onto a 5 mL HisTrap HP column (Cytiva). Protein elution was performed with 50 mM Tris-HCl pH 8.0, 300 mM imidazole and 300 mM NaCl. Protein fractions were loaded onto a gel filtration column (HiLoad 26/60 Superdex 200 prep grade, Cytiva) equilibrated with 50 mM Tris-HCl pH 8.0 and 150 mM NaCl. The fractions with the highest protein concentration were pooled, concentrated, and stored at -80°C.

### Cloning, expression and purification of YejA^Ec^

*yejA*^Ec^ lacking the sequence coding for the signal peptide was cloned into the pEHisTEV vector [56] generating an N-terminal 6His-tag and transformed into *E. coli* BL21 (DE3) for overexpression. A single colony was picked and used to inoculate 100 mL of LB media supplemented with Km. The starter culture was grown at 37°C with 220 rpm shaking overnight. 10 mL of the overnight starter culture was used to inoculate 1 L of LB media supplemented with Km. The culture was incubated at 37°C with shaking at 220 rpm. Cells grown to an OD_600nm_∼0.6 were induced by the addition of IPTG to a final concentration of 0.5 mM. Cell growth continued at 22°C for 18 hours. Cells were harvested by centrifugation at 6,000 g for 10 minutes and stored at -80°C until further use.

The whole-cell pellet was resuspended in PBS buffer supplemented with 20 mM imidazole (pH 7.5), 3 mM MgCl_2_, deoxyribonuclease (50 unit·mL^-1^), lysozyme (0.1 mg·mL^-1^) and PefaBloc (0.1 mg·mL^-1^). Cell-free lysate was prepared by sonication (10 cycles; 10-second sonication and 10-second interval per cycle) followed by centrifugation (200,000 g for 1 hour) and filtration (0.22 μm). Sonication was performed on ice and the following purification steps were performed at 4 °C. For the IMAC step, the supernatant was loaded on a His-Trap column pre-equilibrated with PBS containing 20 mM imidazole (pH 7.5). The column was washed with 20 column volumes (CVs) of PBS containing a linear imidazole concentration from 20 mM to 250 mM (pH 7.5). The ÄKTAxpress (GE Healthcare) system was used to monitor the UV_280nm_ absorbance of the flow-through. The His_6_-tagged YejA protein was eluted with PBS buffer containing 125 mM imidazole. The purity of different fractions was assessed by SDS-PAGE. Fractions with high purity were collected and TEV protease was added with a protein-to-TEV ratio of 10:1 to cleave the 6His-tag. The sample was dialyzed overnight in 2 L of buffer A (20 mM Tris-HCl pH 7.5, 150 mM NaCl).

The overnight cleaved sample was loaded on a His-Trap column pre-equilibrated with buffer A (reverse-IMAC). The flow-through and 2 CV of buffer A wash were collected and concentrated to a final volume of 500 μL, and injected on a Superdex 200 10/300 GL column, pre-equilibrated with buffer A. Fractions containing YejA protein were collected and concentrated to 10 mg·mL^-1^ using a 50 kDa MW cut-off concentrator (Thermo Fisher Scientific).

### Crystallization and structure determination of YejA^Sm^

Crystallization conditions for YejA^Sm^ at 14 mg·mL^-1^ were screened using QIAGEN kits (Valencia, CA) with a Mosquito nanodrop robot (SPT Labtech). YejA^Sm^ crystals were manually optimized in the condition specified in **Table S3**. Crystals were transferred to a cryo-protectant solution (mother liquor supplemented with 25% PEG 400) and flash-frozen in liquid nitrogen. Diffraction data were collected at 100 K on PROXIMA 2 beamline at SOLEIL synchrotron (Saint-Aubin, France). Data processing was performed using the XDS package [57] (**Table S3**). Because of the diffraction anisotropy, the DEBYE and STARANISO programs developed by Global phasing Ltd were applied to the data scaled with AIMLESS using the STARANISO server (http://staraniso.globalphasing.org). These programs perform an anisotropic cut-off of merge intensity data on the basis of an analysis of local I/σ(I), compute Bayesian estimates of structures amplitudes, taking into account their anisotropic fall-off, and apply an anisotropic correction to the data. The structure was solved by molecular replacement with PHASER [58] using the coordinates of the separate N- and C-terminal lobes of the Cu(I)-methanobactin complex-binding protein MbnE from *M. parvus* OBBP (PDB ID: 5ICQ, [33]) as search models. Inspection of the resulting model using COOT [59] showed strong electron density maps at the lobes interface, which were attributed to peptides likely coming from protein degradation during overexpression. The backbones of the short peptide ligands (2 and 5 amino acid peptides) were modeled at two different places of the interface based on the electron density. Electron density for peptide side chains was more ambiguous and no electron density was present linking the two short bound peptides, indicating that a population of different peptides might be present in the ligand-binding site of YejA^Sm^ molecules within the crystal. Refinement of the structure was performed with BUSTER-2.10 [60] employing TLS groups restraints. Refinement details are shown in **Table S3**. Molecular graphics images were generated using PyMOL (http://www.pymol.org).

### Crystallization and structure determination of YejA^Ec^

Crystallisation trials were carried out using the vapour diffusion sitting drop method on 96-well CrystalQuick plates (Grenier). Purified YejA^Ec^ at a concentration of 10 mg·mL^-1^ (145 μM) was screened against the JCSG-plus, PACT-premier and Index screening plates at 20°C and 4°C. The reservoir volume was 100 μL and the sitting drop consisting of 100 nL of YejA and 100 nL of reservoir solution was dispensed by a Mosquito robot. YejA^Ec^ crystallized in the H3 condition of the JCSG-plus screening plate (**Table S3**). Crystallisation conditions were optimised against different pH (± 0.5), temperature (20°C and 4°C) as well as different protein (5 mg·mL^-1^, 10 mg·mL^-1^ and 30 mg·mL^-1^) and PEG concentrations (± 4% (w/v) with 2% (w/v) interval), on CrystalQuick plates. Microseeding was carried out using a contemporary method with the Seed Bead (Hampton Research). Briefly, following the addition of 10 μL of reservoir solution, the macrocrystals were crushed into microcrystals with the Crystal Crusher (Hampton Research) and transferred to the Seed Bead placed on ice. This step was repeated five times until all of the crushed crystals had been transferred to the Seed Bead. This resulted in ∼50 μL of solution containing crushed crystals in the Seed Bead. The Seed Bead was vortexed on ice (three cycles; 30-second vortex plus 30-second interval per cycle). The sitting drop was made of 100 nL of YejA, 30 nL of the prepared microseed and 70 nL of reservoir solution.

YejA^Ec^ crystals were cryoprotected by transferring to the crystallization buffer supplemented with 20% glycerol (v/v) and flash-cooled in liquid nitrogen for data collection. X-ray data were collected on the I24 beamline at the Diamond Light Source. Data were indexed using Xia2 [61] (**Table S3**). Initial phases were obtained by PHASER [58] molecular replacement using for the N-terminal domain MbnE from *M. parvus* (PDB ID: 5ICQ) and for the C-terminal domain AppA from *B. subtilis* (PBI ID: 4ONY) as search models. An initial model was built by Buccaneer autobuild [62]. Iterative cycles of model building and refinement were completed using phenix.refine [63] and COOT [59]. Similarly to YejA^Sm^ structure, strong electron density was observed at the interface between lobes of YejA^Ec^, which enabled the model building of the backbone of a polypeptide that consists of 12 amino acids; the peptide sequence appears to be a degradation product of YejA, likely formed during overexpression. Water molecules were automatically added by ARP/wARP [64] and further manually checked in COOT. Refinement details are shown in **Table S3**.

## Supporting information

Manuscript File

## DATA AVAILABILITY

Sequencing data have been deposited in SRA (BioProject ID PRJNA760523). The YejA^Sm^ and YejA^Ec^ structure factors and coordinates have been deposited to the Protein Data Bank with PDB IDs 7Z8E and 7Z6F respectively.

## AUTHOR CONTRIBUTIONS

D.T., K.S., and P.M. designed the study and planned the experiments. D.T., S.D., R.J., J.L., and P.M., performed cloning, strain construction, *in vivo* competition experiments, and strain sensitivity testing. D.T. and A.V. performed purification and crystallization of YejA^Sm^, F.Q. and K.B. performed purification and crystallization of YejA^Ec^. S.M., F.Q., and K.B. processed and analyzed crystallographic data. S. I.-I. did *in vitro* liposome transport assays with PHZ and BLM. D.S. assembled and analyzed NGS data. D.T., K.S., and P.M wrote the manuscript with input from K.B. and S.M. D.T., S.M., P.M., and A.V. prepared the figures. All authors provided critical feedback and helped to shape the manuscript.

## ACKNOWLEDGMENTS

The work was supported by a grant from the Ministry of Science and Higher Education of the Russian Federation (agreement No. 075-10-2021-114 from 11 October 2021 to K.S.) and RSF grant No. 19-14-00266 to S.D., D.T. was supported by Russian Foundation for Basic Research grant No. 20-34-90098 and the “Ostrogradski” fellowship from the Embassy of France. The work in the P.M. laboratory was supported by Saclay Plant Sciences-SPS and grant ANR-17-CE20-0011 from the Agence Nationale de la Recherche. K.B. was funded by a Medical Research Council grant (MR/N020103/1). S. I.-I. is supported by the Japan Society for the Promotion of Science Overseas Fellowship. F.Q. was supported by the Chinese Scholarship Council Scheme. J.L. and R.J. were supported by PhD fellowships from the French Ministry of Higher Education, Research and Innovation. This work further benefited from the I2BC crystallization platforms supported by FRISBI ANR-10-INSB-05-01. We acknowledge SOLEIL for the provision of synchrotron radiation facilities (proposal ID 20170872) in using PROXIMA 2 beamline and we thank the staff for assistance in using the beamline. We also acknowledge the Diamond Light Source synchrotron for beamtime access. NGS library preparation and sequencing were performed at Skoltech Sequencing Core Facilities (Dr. Maria Logacheva). We would also like to acknowledge M. Towrie (STFC) for providing access to the Cary Eclipse Fluorescence Spectrophotometer. We thank Dr. Emanuele Biondi, Dr. Quentin Barriere, Dr. Nicolas Busset, Dr. Marina Serebryakova, Dr. Dmitry Ghilarov, and Ignat Gorelov for their valuable help and advice and Dr. Annette Vergunst for the gift of plasmid pIN72.

## LEGENDS FOR SUPPLEMENTARY FIGURES AND TABLES

**Supplementary Figure 1** | CFU assay of *wt* Sm1021, single and double mutants in *yejABEF* and *bacA* genes on the YEB agar plates without PHZ and with 8 μM PHZ. Small black asterisks indicate occasional splashes of mut #1 0 and 10^−1^ dilutions.

**Supplementary Figure 2** | **(A)** The mutations identified by Sanger sequencing in the *bacA* gene of PHZ-resistant mutants selected using the Sm1021 Ω*yejA* strain. The numbers indicate the nucleotide position in the gene, the primers used for PCR amplification and sequencing are shown as blue arrows. (**B)** A fragment of amino acid sequence alignment of BacA (*S. meliloti*) and SbmA (*E. coli*). (**C)** The structure of *E. coli* SbmA dimer (subunits are shown in blue and light blue, PDB ID: 7P34 [10]), the amino acids homologous to those undergoing substitutions in PHZ-resistant mutants (clones B1 and B2) are shown in stick and colored in red.

**Supplementary Figure 3** | Construction and phenotype verification of the *phzD* mutant of *Rhizobium* sp. Pop5 (**A)** Biosynthetic gene cluster of PHZ (*phzEACBD*) in the genome of *Rhizobium* sp. Pop5. A region, internal to *phzD*, cloned into pVO155 for subsequent plasmid insertion mutagenesis is shown. Oligonucleotide primer pairs used for insertion verification are shown as blue and red arrows. (**B)** DNA gel electrophoresis of the PCR products, amplified from the genomic DNA purified from two clones of Ω*phzD* mutant and *wt Rhizobium* sp. Pop5. (**C)** Mass-spectra of whole cells show the loss of the prominent mass-peak corresponding to mature phazolicin (2363.9 [M+H]^+^) in Ω*phzD* mutant in comparison to *wt Rhizobium* sp. Pop5. The induction of *phzD* gene expression from the pSRK plasmid leads to the restoration of PHZ production by the Ω*phzD* mutant. (**D)** *Rhizobium leguminosarum* 4292 growth inhibition zone observed around the *wt Rhizobium* sp. Pop5 colony is not detected for the Ω*phzD* mutant.

**Supplementary Figure 4** | **(A)** Identification by flow cytometry of individual strains, grown in monoculture during the indicated times. Dot plots show the GFP fluorescence in the x-axis and DsRed fluorescence in the y-axis of individual cells in the culture (dots). (**B)** CFU counting for the pure cultures of Sm1021 (up) and Pop5 (down) derivatives on the selective media containing Sm and Tc, respectively. No growth was observed for pure cultures of Sm1021 on Tc10 and Pop5 on Sm500.

**Supplementary Figure 5** | Structure-based sequence alignment between YejA^Sm^ and YejA^Ec^. Their secondary structure is labelled and indicated by coils for α-helices, arrows for β-strands and η for short 3_10_ helices. Identical residues are shown as white letters on a red background. The positions of 11 Yej^Ec^ amino acids (Pro51/Arg49, Leu132/Gln130, Tyr137/Tyr135, Leu162/Asp159, Asp460/Asn447, Arg468/Arg455, Ala480/Arg467, Asn487/Ser474, Glu488/Asp475, Ser503/Tyr488 and Arg504/Tyr489 in YejA^Sm^/YejA^Ec^) involved in dodecapeptide binding are indicated by red triangles.

**Supplementary Figure 6** | **(A)** Sampling sites of the environmentally isolated strains with *phz*-like biosynthetic gene clusters in the genomes. **(B)** Schematic structures of the *phz*-like BGCs found across Proteobacteria. Variants of gene composition found in the genus *Phyllobacterium* (1) and other genera (2) are shown. The proposed functions of the encoded proteins are listed on the right. A multiple alignment of the amino acid sequences of the precursor peptides encoded in the *phz*-like BGCs is shown below. The alignment consensus is shown; color highlighting is based on the chemical properties of the amino acids and their conservation. Note the high degree of conservation for the residues converted into azole cycles and involved in the PHZ binding with the ribosome across the precursors. **(C)** Maximum likelihood phylogenetic tree of PhzD (YcaO cyclodehydratase, key enzyme for azole cycle installation) homologs, built using PhyML (S. Guindon, J.-F. Dufayard, V. Lefort, M. Anisimova, W. Hordijk, and O. Gascuel, Syst. Biol., vol. 59, no. 3, pp. 307–321, Mar. 2010, doi: 10.1093/sysbio/syq010). PhzD sequence from PHZ-producing *Rhizobium* sp. Pop5 is shown in bold.

**Supplementary Table 1** | Bacterial strains and vectors used in the study.

**Supplementary Table 2** | Nucleotide sequences of primers used in the study.

**Supplementary Table 3** | Crystallographic data and refinement parameters.

**Supplementary Table 4** | Isolation sources for the strains with *phz*-like BGCs in the genome.

