## Supplementary material for "The antibiotic phazolicin displays a dual mode of uptake in Gram-negative bacteria": Manuscript File

**SUPPLEMENTARY FIGURES**

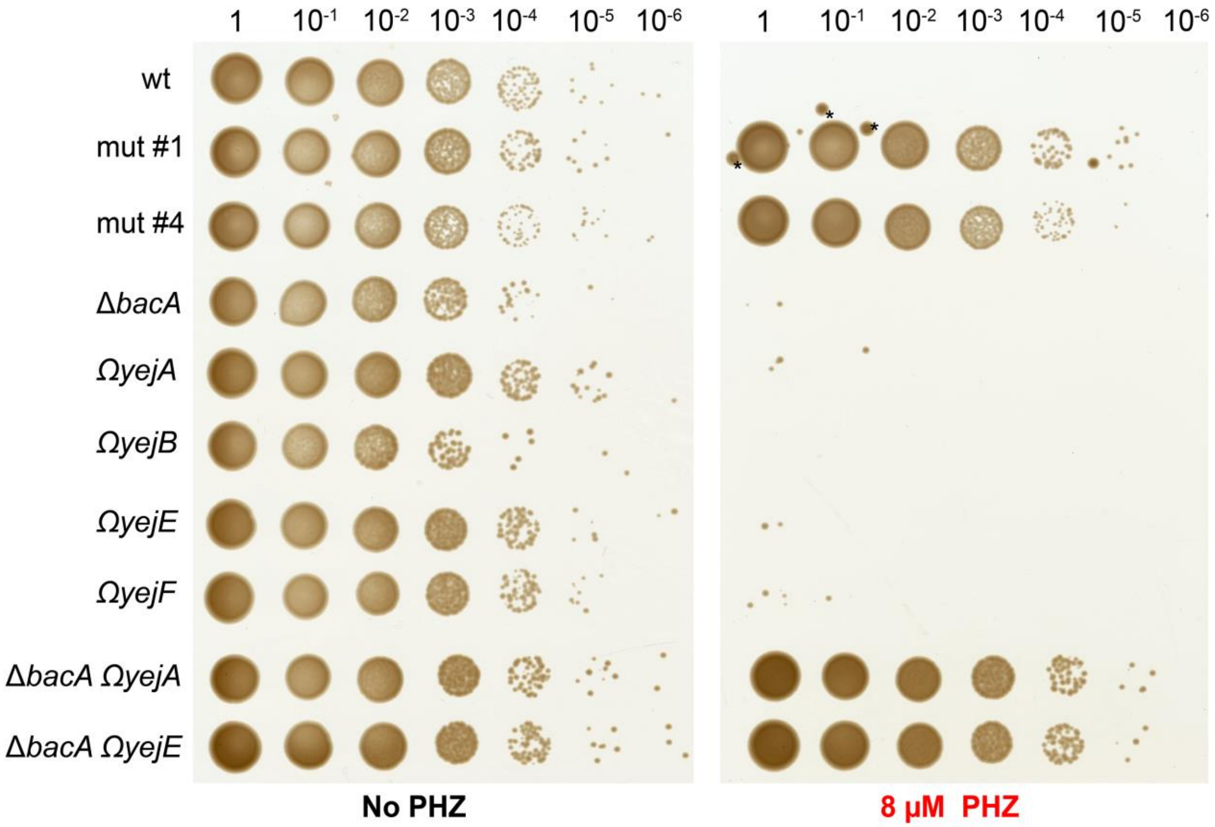


**Supplementary Figure 1 |** CFU assay of *wt* Sm1021, single and double mutants in *yejABEF* and *bacA* genes on the YEB agar plates without PHZ and with 8 µM PHZ. Small black asterisks indicate occasional splashes of mut #1 0 and $10^{-1}$ dilutions.

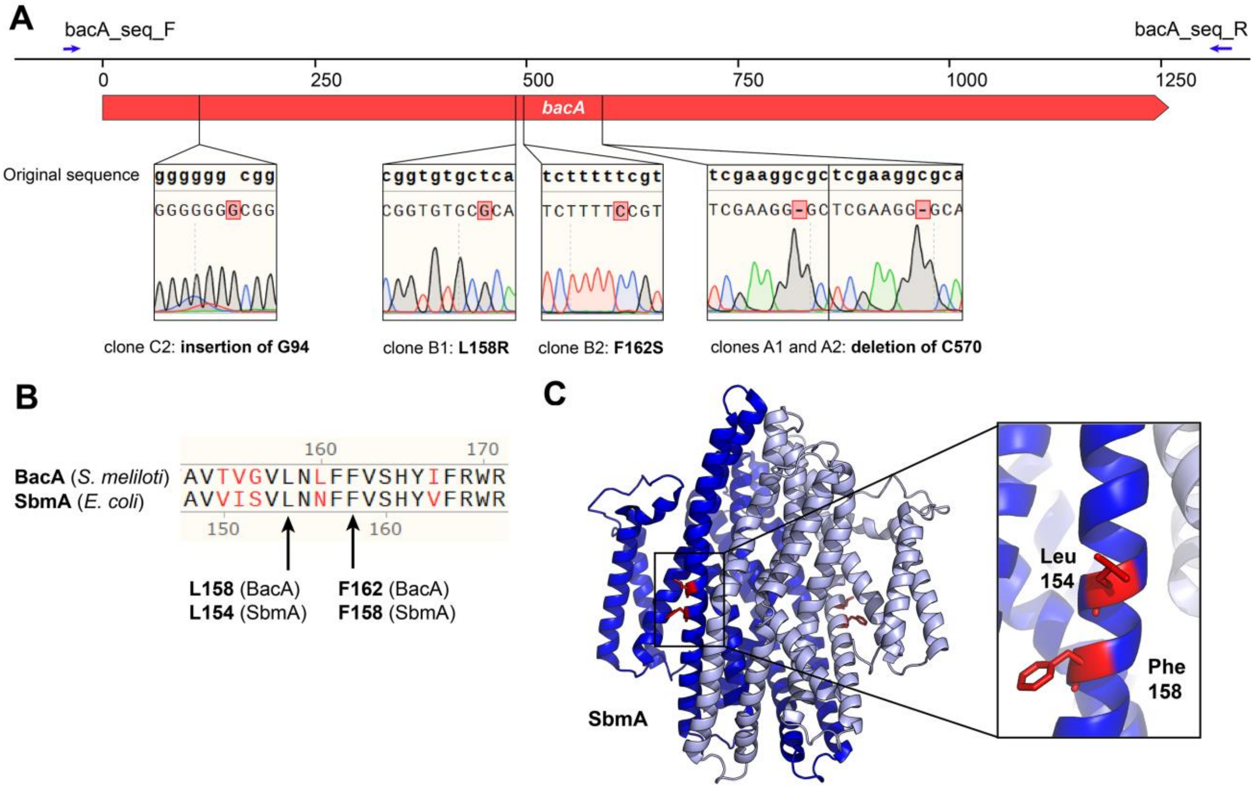


**Supplementary Figure 2 | (A)** The mutations identified by Sanger sequencing in the *bacA* gene of PHZ-resistant mutants selected using the Sm1021 Ω*yejA* strain. The numbers indicate the nucleotide position in the gene, the primers used for PCR amplification and sequencing are shown as blue arrows. **(B)** A fragment of amino acid sequence alignment of BacA (*S. meliloti*) and SbmA (*E. coli*). **(C)** The structure of *E. coli* SbmA dimer (subunits are shown in blue and light blue, PDB ID: 7P34 [10]), the amino acids homologous to those undergoing substitutions in PHZ-resistant mutants (clones B1 and B2) are shown in stick and colored in red.

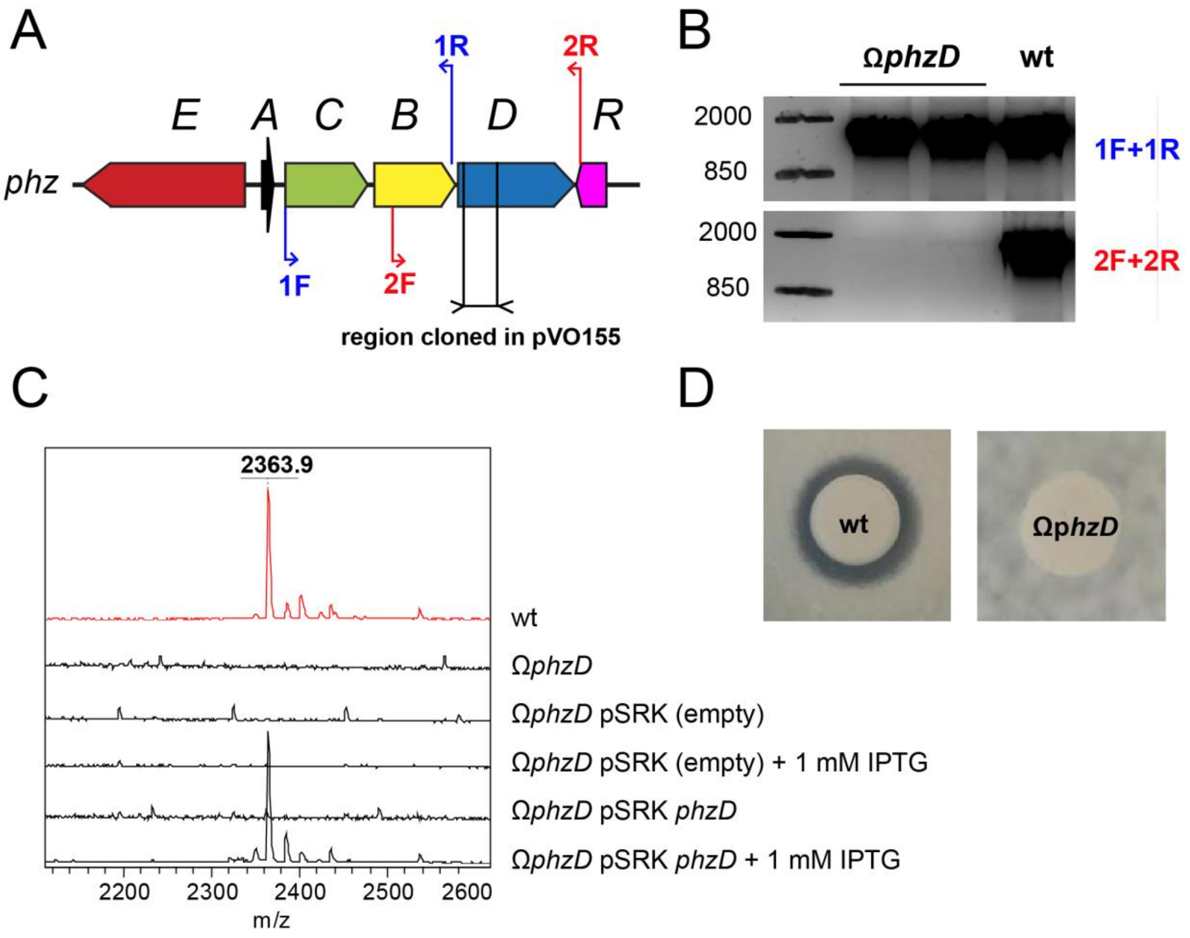


**Supplementary Figure 3 |** Construction and phenotype verification of the *phzD* mutant of *Rhizobium* sp. Pop5 (**A)** Biosynthetic gene cluster of PHZ (*phzEACBD*) in the genome of *Rhizobium* sp. Pop5. A region, internal to *phzD*, cloned into pVO155 for subsequent plasmid insertion mutagenesis is shown. Oligonucleotide primer pairs used for insertion verification are shown as blue and red arrows. **(B)** DNA gel electrophoresis of the PCR products, amplified from the genomic DNA purified from two clones of Ω*phzD* mutant and *wt Rhizobium* sp. Pop5. (**C)** Mass-spectra of whole cells show the loss of the prominent mass-peak corresponding to mature phazolicin (2363.9 $[M+H]^+$) in Ω*phzD* mutant in comparison to *wt Rhizobium* sp. Pop5. The induction of *phzD* gene expression from the pSRK plasmid leads to the restoration of PHZ production by the Ω*phzD* mutant. (**D)** *Rhizobium leguminosarum* 4292 growth inhibition zone observed around the *wt Rhizobium* sp. Pop5 colony is not detected for the Ω*phzD* mutant.

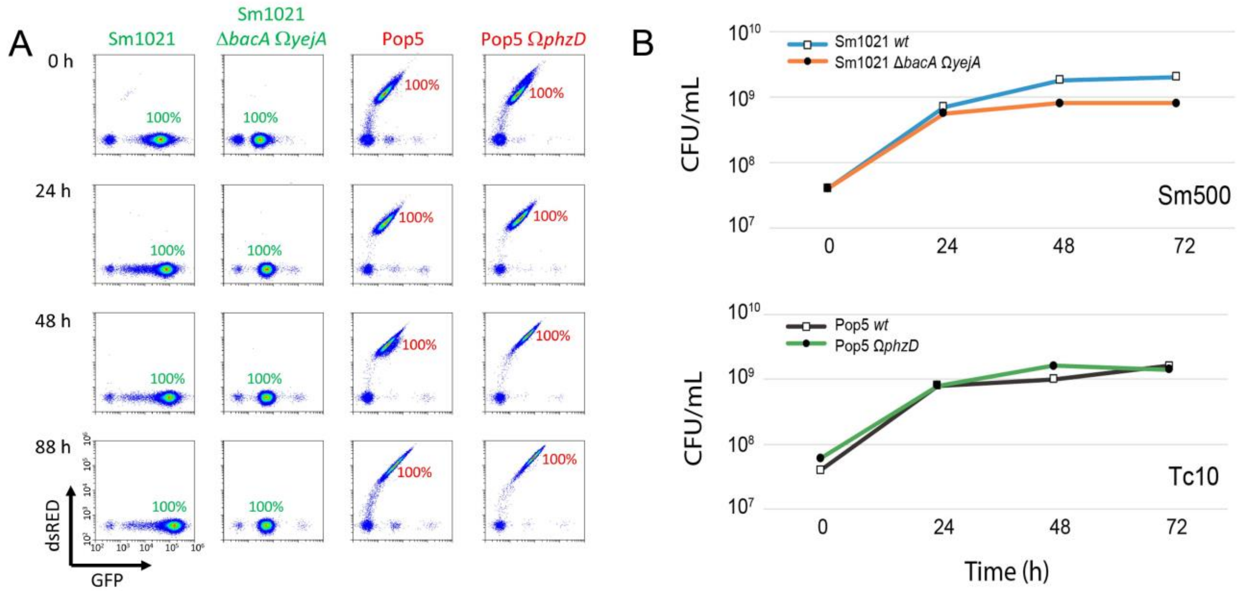


**Supplementary Figure 4 | (A)** Identification by flow cytometry of individual strains, grown in monoculture during the indicated times. Dot plots show the GFP fluorescence in the x-axis and DsRed fluorescence in the y-axis of individual cells in the culture (dots). (**B)** CFU counting for the pure cultures of Sm1021 (up) and Pop5 (down) derivatives on the selective media containing Sm and Tc, respectively. No growth was observed for pure cultures of Sm1021 on Tc10 and Pop5 on Sm500.

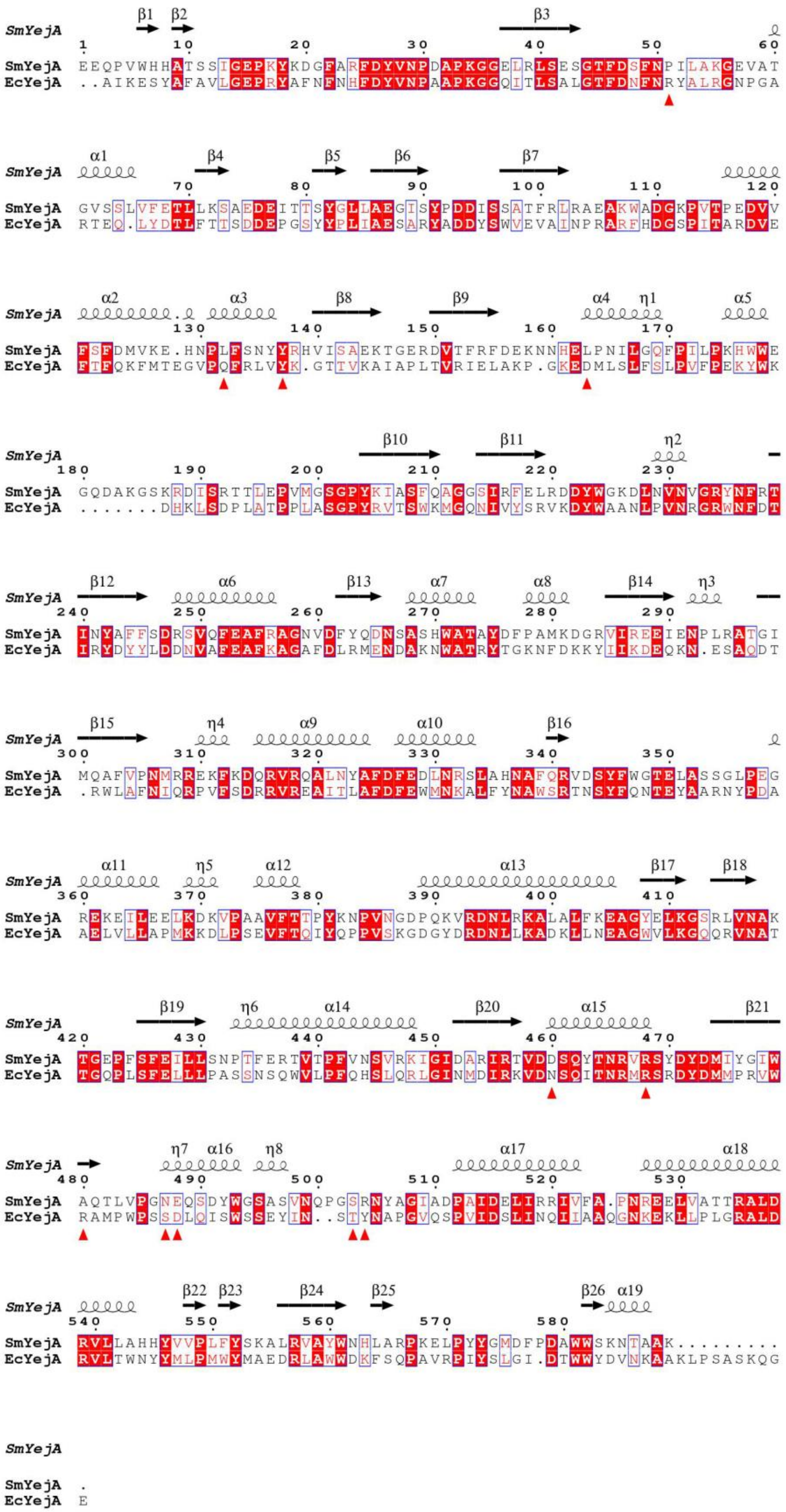

SmYejA EEQPVWHHATSSIGEPKYKDGFARFDYVNPDAPKGGELRLSESGTFDSFNPILAKGEVAT
EcYejA ..AIKESYAFAVLGEPRYAFNFNHFDYVNPAAPKGGQITLSALGTFDNFNRYALRGNPGA
SmYejA GVSSLVFETLLKSAEDEITTSYGLLAEGISYPDDISSATFRLRAEAKWADGKPVTPEDVV
EcYejA RTEQ.LYDTLFTTSDDEPGSYYPLIAESARYADDYSWVEVAINPRARFHDGSPITARDVE
SmYejA FSFDMVKE.HNPLFSNYYRHVISAEKTGERDVTFRFDEKNNHELPNILGQFPILPKHWWE
EcYejA FTFQKFMTEGVPQFRLVYK.GTTVKAIAPLTVRIELAKP.GKEDMLSLFSLPVFPEKYWK
SmYejA GQDAKGSKRDISRTTLEPVMGSGPYKIASFQAGGSIRFELRDDYWGKDLNVNVGRYNFRT
EcYejA .......DHKLSDPLATPPLASGPYRVTSWKMGQNIVYSRVKDYWAANLPVNRGRWNFDT
SmYejA INYAFFSDRSVQFEAFRAGNVDFYQDNSASHWATAYDFPAMKDGRVIREEIENPLRATGI
EcYejA IRYDYYLDDNVAFEAFKAGAFDLRMENDAKNWATRYTGKNFDKKYIIKDEQKN.ESAQDT
SmYejA MQAFVPNMRREKFKDQRVRQALNYAFDFEDLNRSLAHNAFQRVDSYFWGTELASSGLPEG
EcYejA .RWLAFNIQRPVFSDRRVREAITLAFDFEWMNKALFYNAWSRTNSYFQNTEYAARNYPDA
SmYejA REKEILEELKDKVPAAVFTTPYKNPVNGDPQKVRDNLRKALALFKEAGYELKGSRLVNAK
EcYejA AELVLLAPMKKDLPSEVFTQIYQPPVSKGDGYDRDNLLKADKLLNEAGWVLKGQQRVNAT
SmYejA TGEPFSFEILLSNPTFERTVTPFVNSVRKIGIDARIRTVDDSQYTNRVRSYDYDMIYGIW
EcYejA TGQPLSFELLLPASSNSQWVLPFQHSLQRLGINMDIRKVDNSQITNRMRSRDYDMMPRVW
SmYejA AQTLVPGNEQSDYWGSASVNQPGSRNYAGIADPAIDELIRRIVFA.PNREELVATTRALD
EcYejA RAMPWPSSDLQISWSSEYIN..STYNAPGVQSPVIDSLINQIIAAQGNKEKLLPLGRALD
SmYejA RVLLAHHYVVPLFYSKALRVAYWNHLARPKELPYYGMDFPDAWWSKNTAAK.........
EcYejA RVLTWNYYMLPMWYMAEDRLAWWDKFSQPAVRPIYSLGI.DTWWYDVNKAAKLPSASKQG
SmYejA .
EcYejA E

**Supplementary Figure 5 |** Structure-based sequence alignment between $YejA^{Sm}$ and $YejA^{Ec}$. Their secondary structure is labelled and indicated by coils for α-helices, arrows for β-strands and η for short $3_{10}$ helices. Identical residues are shown as white letters on a red background. The positions of 11 $Yej^{Ec}$ amino acids (Pro51/Arg49, Leu132/Gln130, Tyr137/Tyr135, Leu162/Asp159, Asp460/Asn447, Arg468/Arg455, Ala480/Arg467, Asn487/Ser474, Glu488/Asp475, Ser503/Tyr488 and Arg504/Tyr489 in $YejA^{Sm}$/$YejA^{Ec}$) involved in dodecapeptide binding are indicated by red triangles.

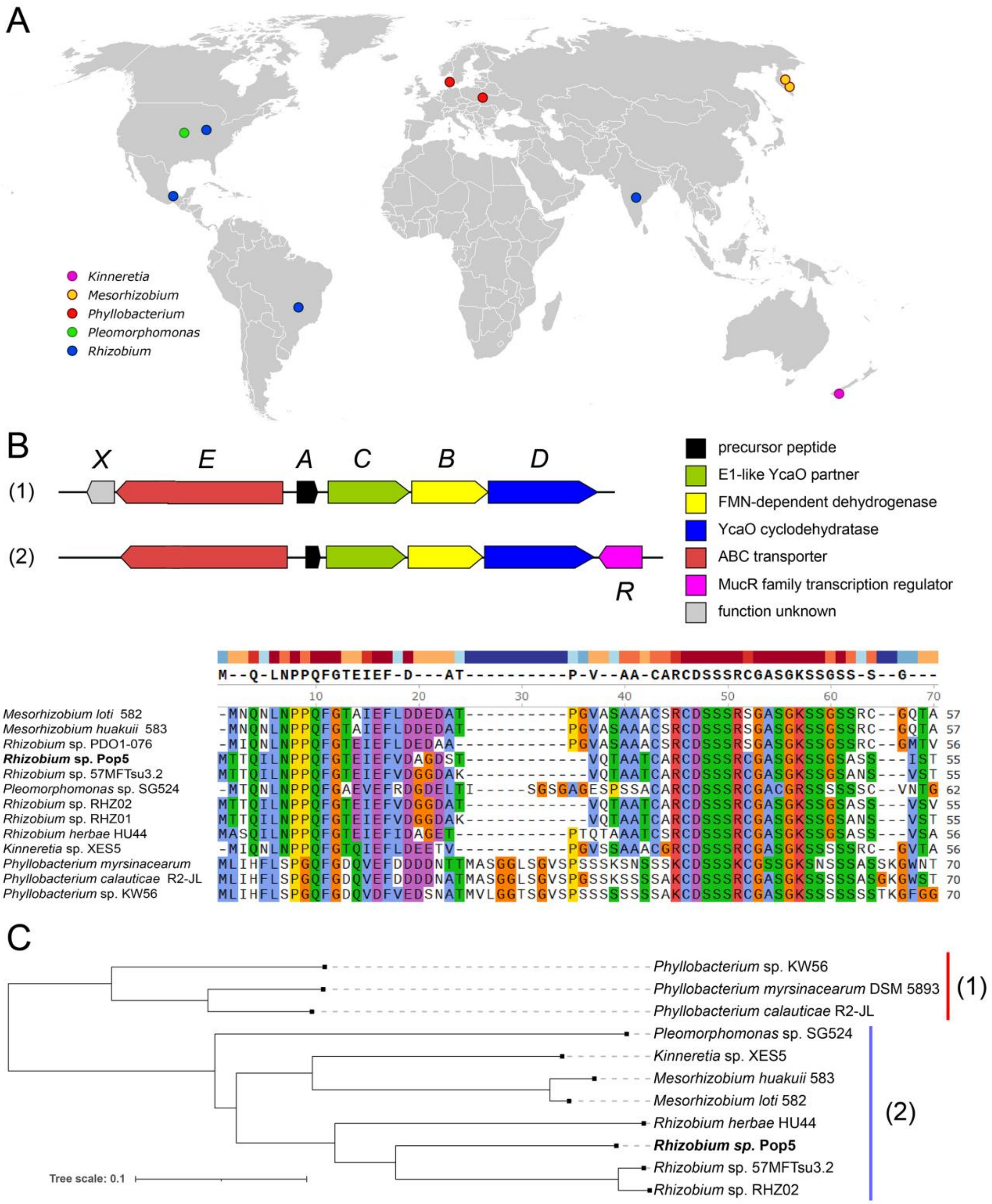


**Supplementary Figure 6 | (A)** Sampling sites of the environmentally isolated strains with *phz*-like biosynthetic gene clusters in the genomes. **(B)** Schematic structures of the *phz*-like BGCs found across Proteobacteria. Variants of gene composition found in the genus *Phyllobacterium* (1) and other genera (2) are shown. The proposed functions of the encoded proteins are listed on the right. A multiple alignment of the amino acid sequences of the precursor peptides encoded in the *phz*-like BGCs is shown below. The alignment consensus is shown; color highlighting is based on the chemical properties of the amino acids and their conservation. Note the high degree of conservation for the residues converted into azole cycles and involved in the PHZ binding with the ribosome across the precursors. **(C)** Maximum likelihood phylogenetic tree of PhzD (YcaO cyclodehydratase, key enzyme for azole cycle installation) homologs, built using PhyML (S. Guindon, J.-F. Dufayard, V. Lefort, M. Anisimova, W. Hordijk, and O. Gascuel, Syst. Biol., vol. 59, no. 3, pp. 307–321, Mar. 2010, doi: 10.1093/sysbio/syq010). PhzD sequence from PHZ-producing *Rhizobium* sp. Pop5 is shown in bold.

#### SUPPLEMENTARY TABLES

**Supplementary Table 1 |** Bacterial strains and vectors used in the study.

**Supplementary Table 2 |** Nucleotide sequences of primers used in the study.

**Supplementary Table 3 |** Crystallographic data and refinement parameters.

**Supplementary Table 4 |** Isolation sources for the strains with *phz*-like BGCs in the genome.

**Supplementary Table 1 | Bacterial strains and vectors used in the study.**

| Strain | Resistance | Description | Reference or source |
|---|---|---|---|
| *S. meliloti* Sm1021 | $Sm^R$ | *Sinorhizobium meliloti* wild-type strain | Common laboratory strain |
| *S. meliloti* Sm1021 Δ*bacA* | $Sm^R$, $Sp^R$ | Sm1021 *bacA654*::Sp (Δ*bacA* null mutant) | [1] |
| *S. meliloti* Sm1021 Ω*yejA* | $Sm^R$, $Km^R$ | *yejA* plasmid insertion mutant | [2] |
| *S. meliloti* Sm1021 Ω*yejB* | $Sm^R$, $Km^R$ | *yejB* plasmid insertion mutant | This study |
| *S. meliloti* Sm1021 Ω*yejE* | $Sm^R$, $Km^R$ | *yejE* plasmid insertion mutant | [2] |
| *S. meliloti* Sm1021 Ω*yejF* | $Sm^R$, $Km^R$ | *yejF* plasmid insertion mutant | [2] |
| *S. meliloti* Sm1021 Ω*tolC* | $Sm^R$, $Km^R$ | *tolC* plasmid insertion mutant | [3] |
| *S. meliloti* Sm1021 Δ*bacA* Ω*yejA* | $Sm^R$, $Km^R$, $Sp^R$ | Double mutant in *yejA* and *bacA* obtained through ϕM12 phage transduction | This study |
| *S. meliloti* Sm1021 Δ*bacA* Ω*yejE* | $Sm^R$, $Km^R$, $Sp^R$ | Double mutant in *yejE* and *bacA* obtained through ϕM12 phage transduction | This study |
| *S. meliloti* Sm1021 mut #1 – #4 | $Sm^R$, $Km^R$ | Phazolicin-resistant mutants selected using transposon library screening | This study |
| *Rhizobium* sp. Pop5 | - | Phazolicin-producing strain (natural isolate) | [4] |
| *Rhizobium* sp. Pop5 Ω*phzD* | $Km^R$ | Mutant with pVO plasmid insertion in *phzD* gene, unable to produce mature phazolicin | This study |
| *Rhizobium leguminosarum* 4292 | $Rif^R$ | PHZ-sensitive strain | Common laboratory strain |
| *E. coli* Rosetta 2 (DE3) pLysS | $Cm^R$ | The strain used for heterologous protein expression | Novagen |
| *E. coli* BL21 (DE3) | - | The strain used for heterologous protein expression | Novagen |
| *E. coli* DH5α | - | The strain used for molecular cloning of all constructs | Common laboratory strain |
| *E. coli* MFDpir Δ*dapA* | - | Donor strain used for transposon library creation, auxotroph for DAP synthesis | [5] |
| *E. coli* BW25113 Δ*tolC* | $Km^R$ | Keio collection strain with the deletion of *tolC* gene encoding the major outer membrane efflux pump | [6] |
| **Vector** | **Resistance** | **Description** | **Reference or source** |
| pDG71 | $Tc^R$ | Constitutive production of GFP(mut3) under *trp* promoter | [7] |
| pIN72 | $Tc^R$ | Constitutive production of the DsRed fluorescent protein | [8] |
| pVO155*nptII**gfp* | $Ap^R$ | Insertional gene inactivation in rhizobia | [9] |
| pET29b(+) | $Km^R$ | Expression vector for C-terminally 6His-tagged proteins in *E. coli* | Novagen |
| pET29 *yejA* (Sm) CHis6 | $Km^R$ | Expression of C-terminally 6His-tagged $YejA^{Sm}$ in *E. coli* | This study |
| pEHisTEV | $Km^R$ | Expression vector for TEV protease cleavable N-terminal 6His-tag fusions in *E. coli* | [10] |
| pEHisTEV *yejA* (Ec) NHis6 | $Km^R$ | Expression of TEV protease cleavable N-terminal 6His-tag-$YejA^{Ec}$ fusion in *E. coli* | This study |

| | | | |
|---|---|---|---|
| pSAM_Ec | $Ap^R$ | Plasmid for *Mariner* transposon library creation | [11] |
| pRK600 | $Cm^R$ | Helper plasmid for conjugal DNA transfer | [12] |
| pSRK | $Gm^R$ | Broad-host range vector for inducible (*lac* promoter) protein expression, oriV (pBBR5 derivative) | [13] |
| pSRK *phzD* | $Gm^R$ | *phzD* from *Rhizobium* sp. Pop5 | This study |
| pSRK *bacA* (Sm) | $Gm^R$ | *bacA* from *Sinorhizobium meliloti* Sm1021 | This study |
| pSRK *bclA* (Bsp) | $Gm^R$ | *bclA* from *Bradyrhizobium* sp. ORS 285 | This study |
| pSRK *sbmA* (Ec) | $Gm^R$ | *sbmA* from *Escherichia coli* MG1655 | This study |
| pSRK *bacA* (Ba) | $Gm^R$ | *bacA* from *Brucella abortus* 2308 | This study |
| pSRK *yejABEF* (Sm) | $Gm^R$ | *yejABEF* from *Sinorhizobium meliloti* Sm1021 | This study |
| pSRK *yejABEF* (Ec) | $Gm^R$ | *yejABEF* from *Escherichia coli* MG1655 | This study |
| pSRK *$nppA_1A_2BCD$* (Pa) | $Gm^R$ | *$nppA_1A_2BCD$* from *Pseudomonas aeruginosa* PA14 | This study |

Ap – ampicillin, Cm – chloramphenicol, Gm – gentamycin, Km – kanamycin, Rif – rifampicin, Sm – streptomycin, Sp – spectinomycin, Tc – tetracycline

**Supplementary Table 2 | Nucleotide sequences of primers used in the study.**

| Primer name | Primer sequence (5'-3') | Purpose |
|---|---|---|
| bacA_Sm_pSRK_GA_F | ataacaatttcacacaggaaacagcatatgttccaatccttcttcccc | Molecular cloning of *bacA (S. meliloti* Sm1021*), bacA (B. abortus), bclA (Bradyrhizobium* sp. ORS285*)* and *sbmA (E. coli* MG1655*)* genes into the pSRK plasmid by Gibson Assembly (*bacA*$^{Sm}$) or restriction ligation protocol (others). |
| bacA_Sm_pSRK_GA_R | cgaggtcgacggtatcgatattacagagccagctcttcc | |
| bacA_Ba_NdeI_F | attataCATATGtttgcgtcatttttccccg | |
| bacA_Ba_XbaI_R | atattaTCTAGAtcagctcgcccctggttc | |
| bclA_Bsp_NdeI_F | attattaCATATGaacaatttgcgctcgaccc | |
| bclA_Bsp_XbaI_R | attataaTCTAGActactcggcgccacccgcc | |
| sbmA_Ec_NdeI_F | attattaCATATGtttaagtctttttcccaaagc | |
| sbmA_Ec_XbaI_R | atattaTCTAGAttagctcaaggtatgggttacttc | |
| pSRK_GA_F | atgctgtttcctgtgtgaaattg | |
| pSRK_GA_R | tatcgataccgtcgacctcg | |
| yejA_Sm_pSRK_GA_F | ataacaatttcacacaggaaacagcataatgccaaacttctgcaggaccg | Molecular cloning of *yejA (S. meliloti* Sm1021*)* gene into the pSRK plasmid by Gibson Assembly. |
| yejA_Sm_pSRK_GA_R | cgaggtcgacggtatcgatatcattttgcagccgtgtttttcg | |
| yejA_Ec_NdeI_F | taatattaaCATATGattgtgcgcatactgc | Molecular cloning of *yejABEF* genes (*E. coli* MG1655) into the pSRK plasmid. |
| yejF_Ec_XbaI_R | ataattaTCTAGAtcagctcaacgccagtagctg | |
| yejE_Ec_mut_F | tcatcctgcgtcacatgttgcctaatgccatg | Internal NdeI site elimination from the *yejE*$^{Ec}$ gene. |
| yejE_Ec_mut_R | tggcattaggcaacatgtgacgcaggatgatact | |
| nppA1_Pa_NdeI_F | taatattaaCATATGcgtcgcctctccttc | Molecular cloning of *nppA*$_1$*A*$_2$*BCD* genes (*P. aeruginosa* PA14) into the pSRK plasmid. |
| nppD_Pa_SacI_R | attataaGAGCTCtcagttttccgcgcttgcc | |
| yejA_Sm_NoSP_NdeI_F | atttattaCATATGgaggaacaacccgtctggcacc | Molecular cloning of *yejA* gene (*S. meliloti* Sm1021) into the pET19b plasmid. |
| yejA_Sm_XhoI_R | attaattCTCGAGttttgcagccgtgtttttcgacc | |
| yejA_Ec_NoSP_NcoI_F | catgCCATGGctatcaaggaaagctatgcc | Molecular cloning of *yejA* gene (*E. coli* MG1655) into the pEHisTEV plasmid. |
| yejA_Ec_XhoI_R | ccgCTCGAGctactctccctgtttgctgg | |
| bacA_seq_F | gcatcaggaggcaagtccttg | Amplification of Sm1021 *bacA* gene region for subsequent amplicon Sanger sequencing. |
| bacA_seq_R | gaggcgttgccgattatcgag | |
| phzC_1F | atgttttcggtttccccgttcgtac | Verification of the pVO155 plasmid insertion into the *phzD* gene. |
| phzB_1R | gcattaattctcctccggataggcaaagg | |
| phzB_2F | accatcgagtttcccgatg | |
| phzD_2R | gctctcaagctaaagcaaaataaggc | |
| phzD_pVO_SalI_F | attatatGTCGACcgagatatcgtgtgacccc | Cloning of the *phzD* gene 541 bp-long fragment into pVO155. |
| phzD_pVO_XbaI_R | attattaTCTAGAcgccaagaccttcgatagc | |
| phzD_NdeI_F | atattatCATATGcaacggtcatatcgc | Cloning of *phzD* gene into pSRK plasmid. |
| phzD_HindIII_R | gctgttAAGCTTttatgaaaatggcatgggc | |

**Supplementary Table 3 | Crystallographic data and refinement parameters.**

| | **YejA$^{Sm}$#** | **YejA$^{Ec}$** |
|---|---|---|
| PDB code | 7Z8E | 7Z6F |
| Crystallization conditions | 14% PEG 8K, 0.1 M Tris-HCl pH 8.5 0.2 M $MgCl_2$ | 25% PEG 3350, 0.1 M Bis-Tris pH 5.5 |
| Beamline | SOLEIL-PX2 | DLS-I24 |
| Wavelength (Å) | 0.9793 | 0.9686 |
| *Za* | 1 | 1 |
| Space group<br>Cell parameters (Å,°) | $P2_12_12_1$<br>*a*= 59.9<br>*b*= 73.7<br>*c*= 140.8 | $P2_12_12_1$<br>*a*= 70.5<br>*b*= 77<br>*c*= 103.8 |
| Resolution (Å) | 65.34-1.58 (1.67-1.58) | 62.19-1.65 (1.68-1.65) |
| No. of observed reflections | 765515 (40211) | 293068 (13977) |
| No. of unique reflections | 86259 (4780) | 68508 (3281) |
| Completeness spherical (%) | 100 (100) | 99.6 (95.9) |
| Mean I/σ(I) | 11.3 (0.6) | 7.4 (1.0) |
| Completeness spherical Staraniso (%) | *85.7 (22.6)* | - |
| Completeness ellipsoidal Staraniso (%) | *95.5 (59.7)* | - |
| $R_{merge}$ (%) | *10.2 (164)* | 11.6 (135) |
| $R_{pim}$ (%) | *3.6 (53.8)* | 6.2 (72.8) |
| Mean I/σ(I) after Staraniso | *13.1 (1.4)* | - |
| $CC_{1/2}$ | *0.99 (0.54)* | 0.99 (0.50) |
| $R_{cryst}$ (%) | 16.9 | 18.9 |
| $R_{free}$ (%) | 20.2 | 23.2 |
| rms bond deviation (Å) | 0.01 | 0.006 |
| rms angle deviation (°) | 0.93 | 0.81 |
| Average B (Å$^2$)<br>Protein<br>Peptide 1/2<br>Solvent | <br>25<br>27/29<br>35 | <br>19<br>33<br>30 |
| [a]Clashscore<br>MolProbity score | 0.83<br>0.86 | 3.47<br>1.26 |
| [a]Ramachandran plot (%)<br>Favoured<br>Outliers | <br>98.48<br>0 | <br>97.28<br>0 |

Values for the highest resolution shell are in parentheses
$CC_{1/2}$ = percentage of correlation between intensities from random hall-dataset
[a]Calculated with MolProbity
Numbers in italic account for statistical values after ellipsoidal mask application by Staraniso.
### A dataset collected from a crystal, which diffracted anisotropically to 1.702 Å along *a**, 1.645 Å along *b** and 1.577 Å along *c**

**Supplementary Table 4 | Isolation sources for the strains with *phz*-like BGCs in the genome**

| Strain | Isolation source | Accession number and Ref |
|---|---|---|
| *Kinneretia* sp. XES5 | swab of adult *Xenopus laevis* skin | NZ_CP084752, [1] |
| *Mesorhizobium huakuii* 583 | *Oxytropis kamtschatica* root nodules | NZ_CP050298, [2] |
| *Mesorhizobium loti* 582 | *Oxytropis kamtschatica* root nodules | NZ_CP050294, [2] |
| *Phyllobacterium myrsinacearum* DSM 5893 | NA | NZ_SGXB01000013 |
| *Phyllobacterium calauticae* R2-JL | freshwater sediment | NZ_JAGENB010000002, [3] |
| *Phyllobacterium* sp. KW56 | nodule (species unknown) | NZ_JAIQWW010000038 |
| *Pleomorphomonas* sp. SG524 | *Sorghum bicolor* roots | NZ_JAAOYR010000004, [4] |
| *Rhizobium altiplani* BR 10423 | *Mimosa pudica* nodules | NZ_LNCD01000036, [5] |
| *Rhizobium herbae* HU44 | *Cajanus cajan* root | NZ_JAEUAO010000003 |
| *Rhizobium* sp. Pop5 | *Phaseolus vulgaris* root nodule | NZ_AMCP01000684 |
| *Rhizobium* sp. RHZ01 | soil | NZ_ JACUZZ010000027 |
| *Rhizobium* sp. RHZ02 | soil | NZ_JACUZX010000024 |
| *Rhizobium* sp. PDO1-076 | *Populus deltoides* root material | NZ_AHZC01000156, [6] |
| *Rhizobium* sp. 57MFTsu3.2 | NA | NZ_JDWI01000010 |
